# Importin α/β Promote Kif18B Microtubule Association to Spatially Control Microtubule Destabilization

**DOI:** 10.1101/2022.03.30.486445

**Authors:** Sanjay Shrestha, Stephanie C. Ems-McClung, Mark A. Hazelbaker, Amber L. Yount, Sidney L. Shaw, Claire E. Walczak

## Abstract

Tight regulation of microtubule (MT) dynamics is necessary for proper spindle assembly and chromosome segregation. The MT destabilizing Kinesin-8, Kif18B, controls astral MT dynamics and spindle positioning. Kif18B interacts with importin α/β as well as with the plus-tip tracking protein EB1, but how these associations modulate Kif18B is not known. We mapped the key binding sites on Kif18B, made residue-specific mutations, and assessed their impact on Kif18B function. Blocking EB1 interaction disrupted Kif18B MT plus-end accumulation and inhibited its ability to control MT length on monopolar spindles in cells. Blocking importin α/β interaction disrupted Kif18B localization without affecting aster size. *In vitro*, importin α/β increased Kif18B MT association by increasing the on-rate and decreasing the off-rate from MTs, which stimulated MT destabilization. In contrast, EB1 promoted MT destabilization without increasing lattice binding *in vitro*, which suggests that EB1 and importin α/β have distinct roles in the regulation of Kif18B-mediated MT destabilization. We propose that importin α/β-spatially modulate Kif18B association with MTs to facilitate its MT destabilization activity. Our results suggest that Ran-regulation is important not only to control molecular motor function near chromatin but also provides a spatial control mechanism to modulate MT binding of NLS-containing spindle assembly factors.

## Introduction

The mitotic spindle is a macromolecular machine composed of dynamic microtubules (MTs) and associated proteins responsible for segregation of the chromosomes during mitosis. Proper spindle assembly requires tight spatial and temporal control of MT dynamics and organization that is governed in part by the RanGTP gradient (Kalab and Heald, 2008). RanGTP is high in the vicinity of chromosomes and dissipates steeply toward the spindle poles (Kalab *et al*., 2002; Caudron *et al*., 2005; Kalab *et al*., 2006). RanGTP modulates the activity of multiple spindle assembly factors through the nuclear transport receptors importin α/β, which bind to proteins containing a nuclear localization signal (NLS) (Kalab and Heald, 2008). In regions where RanGTP is high, RanGTP binding to importin β causes a conformational change, which induces importin α to release its cargo, activating the cargo locally within the spindle. Ran-regulated targets include molecular motor proteins that help organize the spindle MT arrays as well as a number of proteins that regulate spindle MT dynamics (Gruss *et al*., 2001; Wiese *et al*., 2001; Trieselmann *et al*., 2003; Ems-McClung *et al*., 2004; Ribbeck *et al*., 2006; Sillje *et al*., 2006; Tahara *et al*., 2008).

Members of both the Kinesin-13 and Kinesin-8 families of molecular motors play critical roles in controlling MT dynamics (Walczak *et al*., 2013). The Kinesin-13 proteins are strictly MT depolymerizing enzymes (Desai *et al*., 1999), whereas the Kinesin-8 proteins are MT plus-end directed motors and MT plus-end destabilizing enzymes (Gupta *et al*., 2006; Varga *et al*., 2006; Hunter *et al*., 2022). The yeast Kinesin-8, Kip3p, can directly depolymerize stabilized MTs in a length dependent manner (Varga *et al*., 2006), albeit less robustly than the Kinesin-13 proteins, while the vertebrate Kinesin-8 proteins, Kif18A, Kif18B, and Kif19A have more subtle effects (Mayr *et al*., 2007; Du *et al*., 2010; Niwa *et al*., 2012; Shin *et al*., 2015; McHugh *et al*., 2018). Kinesin-8 and Kinesin-13 proteins may also act cooperatively to regulate MT dynamics (Tanenbaum *et al*., 2011; McHugh and Welburn, 2023). During mitosis, Kif18A is enriched on kinetochores and is important in controlling chromosome oscillations (Stumpff *et al*., 2008). Analysis of the purified protein suggests that it may have weak MT destabilization activity (Mayr *et al*., 2007) but most likely acts as a capping protein (Du *et al*., 2010). Kif18B localizes to the plus ends of astral MTs (Lee *et al*., 2010) and is important in controlling astral MT dynamics (Walczak *et al*., 2016) and spindle positioning (McHugh *et al*., 2018; Moreci and Lechler, 2021). Full-length Kif18B protein could not depolymerize stabilized MTs, but a truncated derivative lacking the C-terminal tail had weak MT depolymerization activity (McHugh *et al*., 2018).

The mechanisms by which Kif18B modulates MT dynamics are likely complex due to the interaction of its tail domain with multiple binding partners in a spatially regulated manner. Kif18B binds directly to EB1, and its localization to astral MT plus ends is diminished in EB1 knockdown cells (Stout *et al*., 2011; Tanenbaum *et al*., 2011). In contrast, EB1 localization is not affected by Kif18B knockdown (Stout *et al*., 2011; Tanenbaum *et al*., 2011). Consistent with that finding, Kif18B is not uniformly colocalized with EB1, which is found on all MT plus-ends, and is instead enriched on astral MTs near the cortex (Stout *et al*., 2011; Tanenbaum *et al*., 2011; van Heesbeen *et al*., 2017). Super-resolution imaging of mammalian cells showed two spatially resolved localization patterns of Kif18B on astral MTs. While Kif18B and EB1 colocalized on astral MTs closer to the poles, a subset of Kif18B was localized distally to EB1 by 189 nm on astral MTs near the cortex in the vicinity of the metaphase plate (Stout *et al*., 2011; Shin *et al*., 2015). Further evidence of spatial regulation of Kif18B is provided by lifetime imaging of MTs in PtK2 cells depleted of Kif18B. Analysis of EB1 MT plus-tip dynamics in these cells showed that MTs in the polar region near the cortex had reduced dynamics compared to those MTs above and below the spindle (Walczak *et al*., 2016).

In addition to binding EB1, Kif18B has multiple other binding partners. Kif18B physically interacts with MCAK at astral MT ends (Tanenbaum *et al*., 2011; McHugh and Welburn, 2023), where Kif18B and MCAK coordinately modulate MT dynamics. The tail of Kif18B also binds to importin α/β (Stout *et al*., 2011), but whether this binding is simply utilized for nuclear localization or whether Kif18B is another Ran-regulated factor is not known. One possibility is that Kif18B is spatially regulated through the RanGTP network. However, this creates a paradox because the RanGTP gradient is thought to act locally around the chromosomes, yet Kif18B localization is most prominent near the cortex. Kif18B MT destabilization activity is proposed to be important for modulating MT forces between the cortex and spindle poles that impact spindle positioning and spindle bipolarization (van Heesbeen *et al*., 2017; McHugh *et al*., 2018; Moreci and Lechler, 2021). The Kif18B tail also contains a second MT binding site that is important in modulating both motor diffusion and processivity on MTs (Shin *et al*., 2015; McHugh *et al*., 2018). Deletion of the tail disrupts plus-end MT localization in cells (Stout *et al*., 2011; McHugh *et al*., 2018), which is mediated by increases in the off-rate from MTs, reduction in the run length of the motor on the MT, and reduction of the motor dwell time at MT plus ends, despite increased motor velocities (Shin *et al*., 2015; McHugh *et al*., 2018). Previous studies using computer simulations of how a MT destabilizer affects MT dynamics indicate that on-rate, processivity, and off-rate at MT plus ends are critical factors for reducing MT length (Weaver *et al*., 2011). Thus, Kif18B tail-mediated MT and plus-end binding appear to be key for Kif18B function, which we propose is modulated differentially by its interactors, importin α/β and EB1.

In this study we developed a series of inter-molecular FRET reporters to precisely map the individual binding sites of Kif18B with both importin α and EB1. We found that importin α/β and EB1 compete for binding to the Kif18B tail, and that importin α/β increases Kif18B MT destabilization activity by loading Kif18B onto the MT lattice. In contrast, EB1 reduces MT length without increasing overall Kif18B MT binding. Consistent with the effects of EB1 knockdown regulating Kif18B localization in cells, we show that the specific interaction of Kif18B with EB1 is critical for MT plus-end localization and regulation of astral MT length in monopolar spindles. On the other hand, blocking the interaction of Kif18B with importin α/β by mutation of the NLS does not hinder the ability of Kif18B to control MT length in monasters but does prevent bipolar spindle formation when it is overexpressed. Most notably, the NLS mutations compromise enrichment of Kif18B on astral MTs, consistent with importin α/β spatially regulating Kif18B localization. We propose that Ran-regulation is important not only to control motor function near chromatin but that the Ran-system is an overall spatial control mechanism to control MT binding affinity of MT associated proteins.

## Results

### Kif18B Tail Interacts with Importin α/β, EB1, and Microtubules

Kif18B has an N-terminal kinesin catalytic domain, a central stalk region, and a C-terminal tail that has two putative nuclear localization signal (NLS) sequences, one non-canonical bipartite and one monopartite, and three putative EB1 binding domains (EBBD) (Figure 1A) (Lee *et al*., 2010; Stout *et al*., 2011; Tanenbaum *et al*., 2011). To understand how Kif18B interacts with its different binding partners, we designed a series of Förster resonance energy transfer (FRET) reporters to look at protein-protein interactions. Either importin α or EB1 were fused to CyPet as donors, and the Kif18B tail was fused to YPet as the acceptor (Figure 1B). In addition to the full-length tail (aa 605-828), we also generated two tail fragments (aa 605-707 and aa 708-828) to separate the putative NLS and EBBD sites (Figure 1C). To test interactions between the Kif18B tail and importin α, we incubated importin α-CyPet + importin β with increasing molar concentrations of YPet-Kif18B(605-828), YPet-Kif18B(605-707), or YPet-Kif18B(708-828) and measured FRET (Figures 1D and S1A). Importin α-CyPet induced FRET with all three YPet-Kif18B tail fragments, suggesting that the tail of Kif18B contains at least two independent importin α binding domains. Addition of RanQ69L, a mutant form of Ran in which RanGTP cannot be hydrolyzed (Bischoff *et al*., 1994), to importin α-CyPet/importin β and YPet-Kif18B tail proteins disrupted their interaction as seen by a reduction in their FRET ratio, demonstrating Ran regulation of these interactions (Figure S1B). Incubation of EB1-CyPet with increasing molar concentrations of the YPet-Kif18B tail fragments resulted in a dose-dependent increase in FRET with each tail fragment, suggesting that the Kif18B tail also has at least two independent EB1 binding domains (Figures 1E and S1C).

**Figure 1.**
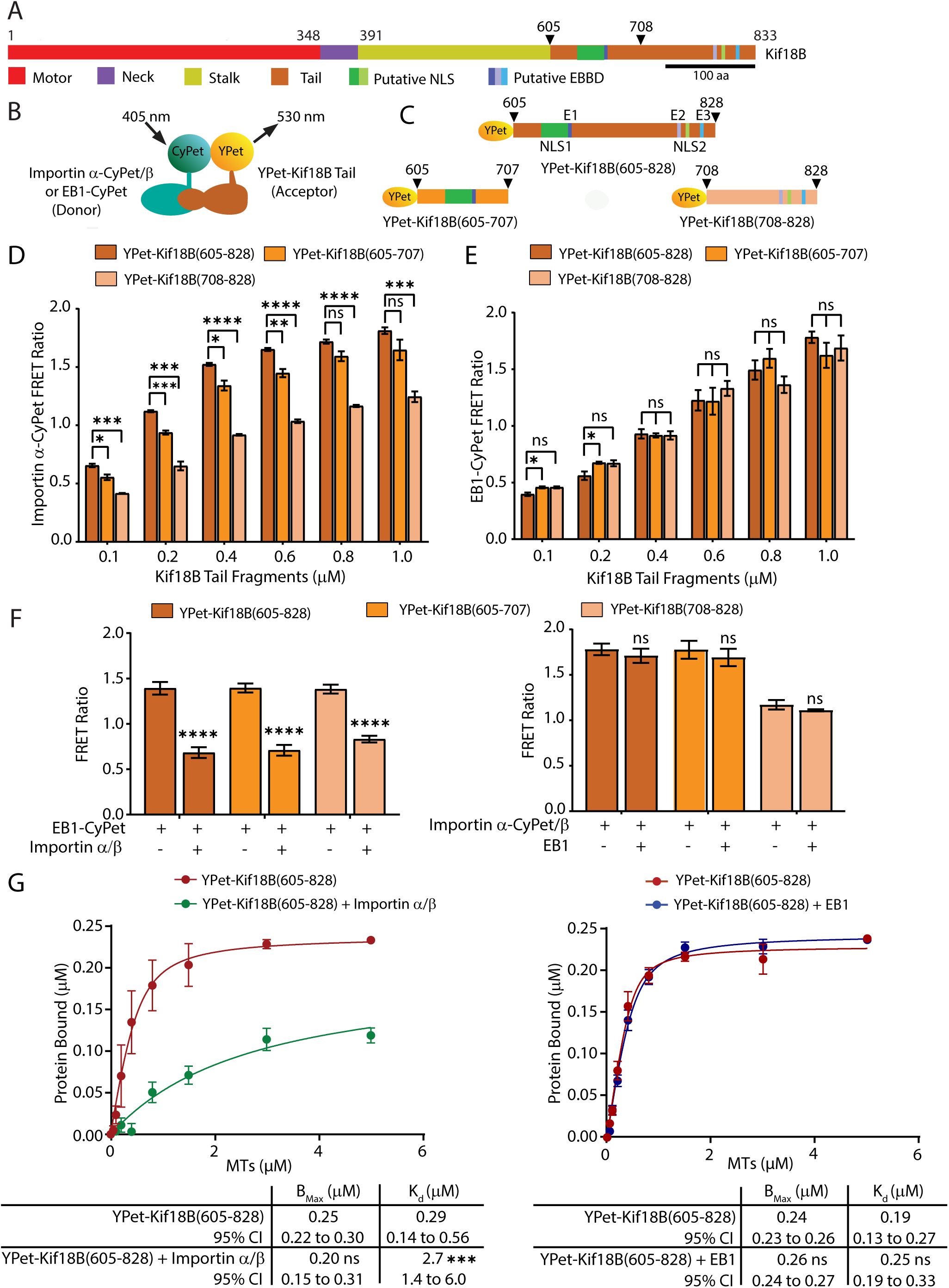
Importin α/β competes with EB1 for binding to Kif18B tail and impairs its microtubule binding. (A) Schematic of full-length human Kif18B showing its domain organization and putative NLS (green) and EB1-binding domains (EBBD) (blue). The arrowheads indicate the beginning residues of the truncated Kif18B proteins used in this study. Scale bar, 100 amino acids (aa). (B) Schematic of FRET between importin α-CyPet/importin β or EB1-CyPet and YPet-Kif18B tail fragments. (C) Schematics of constructs used in this study with the beginning and ending residues of the truncated proteins indicated, and the NLS and EBBD sites marked as in (A). (D and E) Reactions containing 100 nM importin α-CyPet and 400 nM importin β (D) or 100 nM EB1-CyPet (E) were incubated with increasing concentrations of the three different YPet-Kif18B tail proteins and measured for FRET. The normalized FRET ratios at 530 nm are plotted as the mean ± SEM from three independent experiments. (F) Competition experiments with 800 nM YPet-Kif18B tail fragments were performed between 100 nM EB1-CyPet and 400 nM unlabeled importin α/β (left) or between 100 nM importin α-CyPet/importin β and 400 nM unlabeled EB1 (right) in at least three independent experiments. FRET was measured, and the normalized FRET ratios graphed as in (D). (G) YPet-Kif18B(605-828) was incubated with increasing concentrations of MTs in the presence or absence of importin α/β (left) or EB1 (right). The fluorescence of YPet in the supernatant and pellet samples was measured, and the fractional binding of YPet-Kif18B(605-828) in molarity was calculated from at least three independent experiments. The normalized data was plotted as the mean ± SEM and fit to the quadratic binding equation. ns, not significant; *, p<0.05; **, p<0.01; *** p<0.001; ****, p<0.0001.

The putative NLS and EBBD sites are juxtaposed to one another in the tail of Kif18B (Figure 1A). To ask whether the interaction of the Kif18B tail with importin α/β and EB1 is competitive, YPet-Kif18B tail fragments were incubated with EB1-CyPet and a four-fold molar excess of unlabeled importin α/β relative to EB1-CyPet, and the FRET ratio was effectively reduced by 50% (Figure 1F, left), indicating that there is competition for binding between EB1 and importin α/β to the tail of Kif18B. In contrast, when YPet-Kif18B fragments were added to importin α-CyPet/importin β and a four-fold molar excess of unlabeled EB1, EB1 was unable to compete with importin α/β for binding to the tail of Kif18B (Figure 1F, right). Addition of 30X or 40X molar excess of unlabeled EB1 to reactions of YPet-Kif18(605-828) with importin α-CyPet/importin β resulted in only a slight 12-17% decrease in FRET (Figure S1D). Together our data show that the tail of Kif18B contains distinct binding sites that mediate interactions with importin α/β, and EB1, and the Kif18B interaction with importin α/β may have a higher affinity than with EB1.

Members of the Kinesin-8 family have an ATP-independent MT binding domain in the tail that is important for motor binding and processivity, as well as for MT cross-linking (Stumpff *et al*., 2011; Su *et al*., 2011; Weaver *et al*., 2011; McHugh *et al*., 2018). To assess Kif18B tail MT binding, we carried out MT pelleting assays in which YPet-Kif18B(605-828) was incubated with increasing amounts of MTs in the presence or absence of importin α/β or EB1 (Figure 1G). Addition of importin α/β resulted in a mild and non-significant 25% reduction in the maximal binding of the tail to MTs but resulted in a 9-fold increase in the *K*_d_ for MTs (p<0.001) (left); whereas addition of EB1 had no effect on maximal binding nor on the *K*_d_ (Figure 1G, right). These results suggest that importin α/β may modulate Kif18B MT binding through the tail. These results are similar to our previous work with the Kinesin-14 XCTK2, wherein importin α/β regulate the interaction of the XCTK2 tail with MTs to control its ability to cross-link MTs (Ems-McClung *et al*., 2004; Ems-McClung *et al*., 2020), suggesting a common regulatory mechanism for importin-regulated MT binding of kinesin tails.

### Kif18B Contains Two NLS Sites that Mediate Interaction with Importin α/β

To identify the critical residues for binding to importin α/β, we mutated both halves of the bipartite NLS by changing the basic residues to alanine residues (NLS1) or mutated all four basic residues of the monopartite NLS to alanine residues (NLS2) (Figure 2A) and tested them for interaction with importin α/β in the FRET assay (Figure 2B). Mutation of NLS1 in YPet-Kif18B(605-707) and mutation of NLS2 in YPet-Kif18(708-828) significantly reduced the FRET ratio when incubated with importin α-CyPet/importin β. Mutation of both NLS1 and NLS2 (NLS1/2) also significantly reduced the FRET ratio of YPet-Kif18B(605-828) with importin α-CyPet/importin β (Figure 2B) to the same level as mutation of the NLS in each half of the tail, suggesting there are two functional NLS sites in the tail of Kif18B.

**Figure 2.**
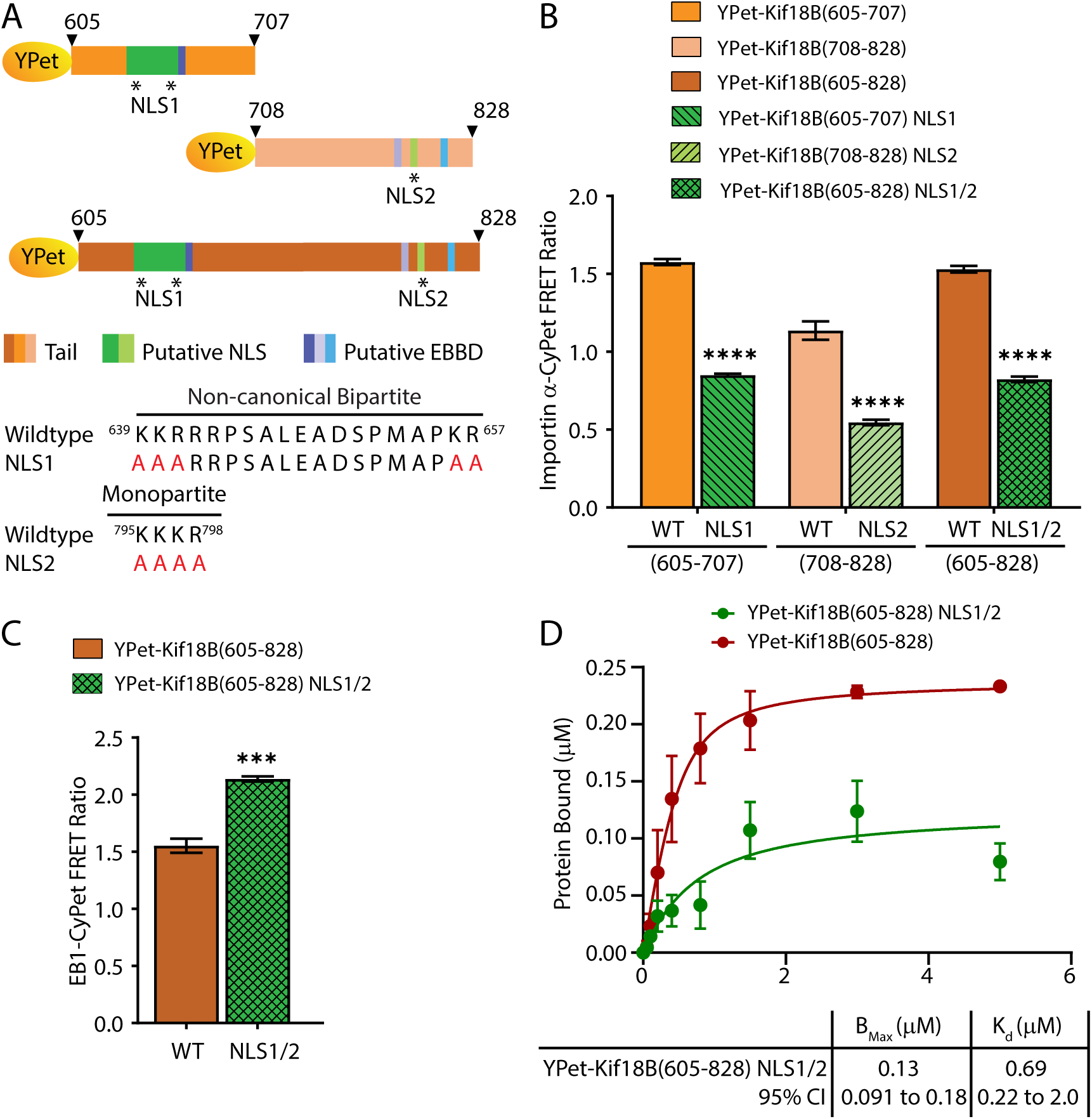
Importin α/β interact with the Kif18B tail through a non-canonical bipartite (NLS1) and a monopartite NLS (NLS2). (A) Schematics of the constructs and mutations (indicated by asterisks) used to identify the critical residues in the Kif18B tail required for importin α/β binding. The wildtype amino acid sequences of Kif18B at the two predicted NLS sites are shown with the mutations introduced at the corresponding residues in red. (B) Normalized FRET ratios following the incubation of 100 nM importin α-CyPet with 400nM importin β and 800 nM wildtype or NLS mutant YPet-Kif18B tail fragments. FRET measurements were performed from three independent experiments and plotted as described in Figure 1D. (C) Normalized FRET ratios of 100 nM EB1-CyPet and 800nM YPet-Kif18B(605-828) or YPet-Kif18B(605-828) NLS1/2. The FRET ratios were measured from three independent experiments and plotted as described in Figure 1D. (D) YPet-Kif18B(605-828) NLS1/2 was incubated with increasing concentrations of MTs and the fractional binding was determined from at least three independent experiments as described in Figure 1G. YPet-Kif18B(605-828) fractional binding from Figxg66
ure 1G (left) is included for comparison. *** p<0.001; **** p<0.0001.

Because of the complex interactions of the Kif18B tail with its different binding partners, we wanted to make sure that mutation of the NLS regions did not impair binding to EB1 or MTs, which would complicate interpretations of functional experiments. The NLS1/2 mutations in YPet-Kif18(605-828) did not impair interaction with EB1-CyPet, rather they slightly increased their interaction (Figure 2C). However, the NLS1/2 mutations caused ∼50% reduction in maximal MT binding and an approximate two-fold increase in the *K*_d_ for MT binding relative to YPet-Kif18B(605-828) (Figures 1G and 2D), suggesting a moderate effect on MT affinity.

### Kif18B Contains Two EB1 Binding Domains

To identify the critical residues for binding to EB1, we mutated the leucine and proline residues (LP) in three predicted EB1 SXIP binding motifs (Honnappa *et al*., 2009) to asparagine (NN) and tested the wild type and mutant constructs for their ability to interact with EB1-CyPet using the FRET assay (Figure 3, A and B). Mutation of E1 in YPet-Kif18B(605-707) and mutation of E3 in YPet-Kif18(708-828) significantly reduced the FRET ratio when incubated with EB1-CyPet (Figure 3B, left), whereas mutation of E2 in YPet-Kif18(708-828) had no effect regardless of the ratio of YPet-Kif18B(708-828) E2 to EB1-CyPet (Figure 3B, right), which suggests that E1 and E3 are functional EB1 binding sites but that E2 is not. Mutation of both E1 and E3 together significantly reduced the FRET ratio of YPet-Kif18B(605-828) with EB1-CyPet to levels similar to either E1 or E3 in their respective half-tail fragments (Figure 3B, left). These results suggest that Kif18B has two separate EB1 binding domains (EBBD) in its tail.

**Figure 3.**
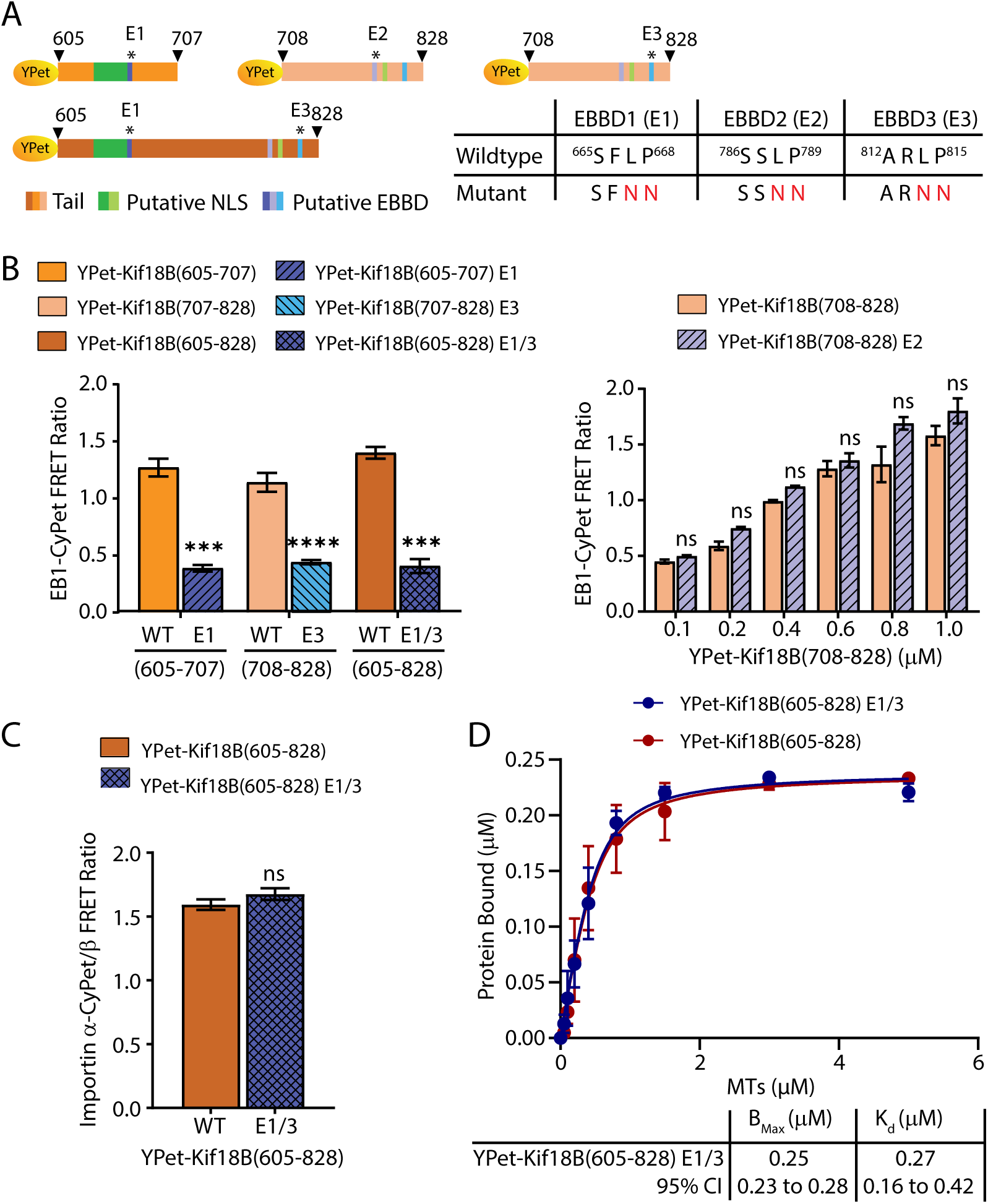
EB1 interacts with Kif18B through two EB1 binding sites in the Kif18B tail. (A) Schematic representations of the constructs and mutations used to map the critical residues needed for EB1 binding where the numbers indicate the residues of Kif18B, and asterisks indicate the mutation sites. The wildtype amino acid sequences of the three putative EB1 binding sites (E1, E2, and E3) and the corresponding mutant sequences are shown with the mutated residues in red. (B) Normalized FRET ratios obtained following incubation of 100 nM EB1-CyPet with 800 nM wildtype or mutant YPet-Kif18B(605-707) E1, YPet-Kif18B(708-828) E3, or YPet-Kif18B(605-828) E1/3 (left). Normalized FRET ratios obtained after incubating 100 nM EB1-CyPet with varying concentrations of wildtype or mutant YPet-Kif18B(708-828) E2 to demonstrate the lack of EB1 binding (right). (C) Normalized FRET ratios of 100 nM importin α-CyPet and 400 nM importin β incubated with 800 nM wildtype or mutant YPet-Kif18B(605-828) E1/3. (B and C) FRET ratios were determined from at least three independent experiments and plotted as in Figure 1D. (D) YPet-Kif18B(605-828) E1/3 was incubated with increasing amounts of MTs to determine fractional binding from at least three independent experiments and was processed as described in Figure 1G. YPet-Kif18B(605-828) fractional binding from Figure 1G (left) is included for comparison. ns, not significant; *** p<0.001; **** p<0.0001.

To test whether mutation of the EBBDs impaired binding to the other partners, we tested the interaction of the EBBD mutated Kif18B(605-828) tail with each of its binding partners (Figure 3, C and D). Mutation of E1/3 in YPet-Kif18(605-828) did not impair interaction with importin α-CyPet/importin β (Figure 3C), nor did it affect its binding to MTs (Figure 3D).

### Interaction with Importin α/β and EB1 are Important for Proper Kif18B Localization on MT Ends and to Modulate Astral MT Dynamics in Cells

To ask whether disruption of the interaction of Kif18B with importin α/β or with EB1 impacts its ability to modulate MT dynamics in cells, we knocked down Kif18B in HeLa cells (Figure S2A), transduced the cells with lentiviral wild-type (WT) GFP-Kif18B or GFP-Kif18B with the NLS1/2 (NLS) or EBBD1/3 (EBBD) mutations, and treated the cells with Monastrol to induce monopolar spindles (Figure 4A). Endogenous Kif18B clearly localized to the tips of astral MTs in control cells (Figure 4A, row 1). This staining was abolished in cells with Kif18B knockdown and resulted in an increase in the MT aster area (Figure 4A, row 2), consistent with previous results (Walczak *et al*., 2016). Lentiviral expressed WT GFP-Kif18B localized to astral MT plus ends like the endogenous Kif18B and reduced the MT aster area (Figure 4A, row 3). Mutation of the NLS in Kif18B reduced MT plus tip localization and reduced MT aster area (Figure 4A, row 4). In contrast, mutation of the EBBD nearly abolished localization of Kif18B to the plus tips of MTs and did not reduce MT aster area (Figure 4A, row 5). When comparing the GFP-Kif18B expression level to MT aster area, we noted that for both the WT and NLS mutant there was a robust negative correlation (Figure S2, B and C) compared to a weaker correlation for the EBBD mutant (Figure S2D). The inability of the EBBD mutant to reduce the aster area and its reduced expression level and correlation with aster area suggests that disruption of the EB1 interaction impacted the ability of Kif18B to destabilize MTs. When analyzing the population of the transduced cells, we noted that the expression levels varied significantly between WT and mutant proteins (Figure S2E). To rule out the possibility that the reduced ability of the GFP-Kif18B EBBD mutant to rescue aster area was due to lower expression levels, we determined the range of expression levels of WT GFP-Kif18B that was able to fully rescue aster area (Figure 4B). We then identified cells transduced with mutant Kif18B NLS and EBBD proteins with a similar range of GFP expression and compared their MT aster area (Figure 4, B and C). Knockdown of Kif18B resulted in a 35% increase in aster area compared to control knockdown with mock expression (Figure 4C). Expression of WT GFP-Kif18B or GFP-Kif18B NLS fully rescued aster area to levels indistinguishable from control knockdown. In contrast, GFP-Kif18B EBBD only partially rescued aster area, which was greater than control knockdown, p<0.0001, but smaller than Kif18B knockdown, p<0.001.

**Figure 4.**
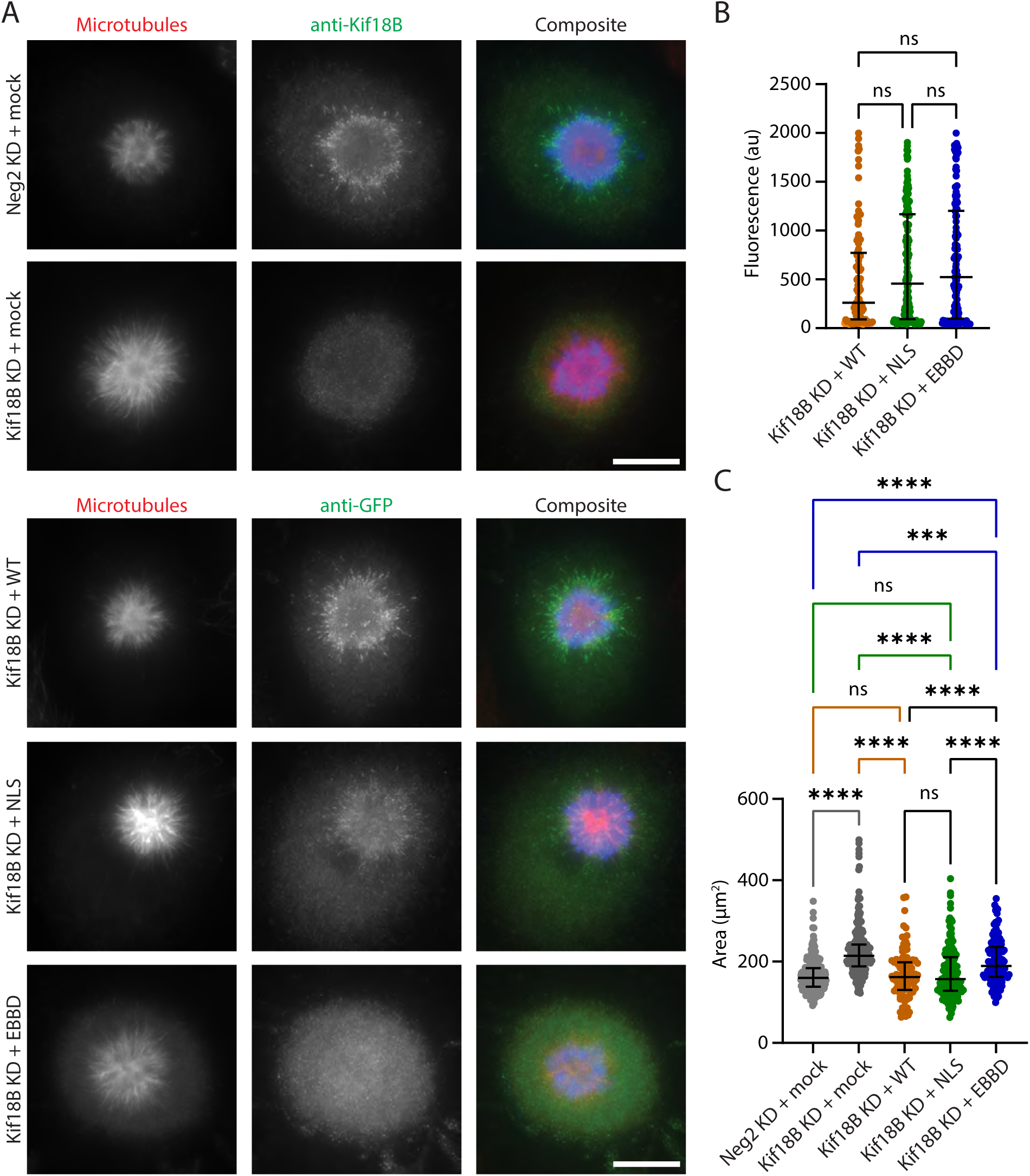
Mutation of the EBBD impacts the ability of Kif18B to control MT length. (A) Cells treated with control Neg2 or Kif18B siRNAs were mock transduced or transduced with WT or mutant lentiviruses, and then fixed and stained for MTs (red), Kif18B (green) or GFP (green), and DNA (blue). Scale bar, 10 µm. (B) The average GFP fluorescence is reported from at least 100 cells from three independent experiments in which each dot represents a single cell, and the median ± interquartile range is indicated. (C) The average MT aster area is reported from the cells in (B) with the median ± interquartile range indicated. ns, not significant; ***, p<0.001; ****, p<0.0001.

We were surprised to observe that there was no change in aster area in cells expressing GFP-Kif18B NLS, as this mutation appeared to compromise the localization of Kif18B. To examine this localization in more detail, we used 3D-SIM super-resolution microscopy to examine the localization of WT GFP-Kif18B versus GFP-Kif18B NLS relative to EB1 on MTs of monopolar spindles (Figure 5). Whereas WT GFP-Kif18B was highly enriched on MT plus ends toward the periphery of the asters, the localization of GFP-Kif18B NLS often appeared more uniformly distributed (Figure 5A). To quantify these differences in localization, we used IMARIS to identify the EB1 and GFP foci and analyzed the number of foci, their surface areas, and the extent of colocalization. There were no differences in the total number of EB1 foci or EB1 surface areas in cells expressing either WT GFP-Kif18B or the NLS mutant (Figure 5, B and C). The number of GFP foci were not different in cells expressing WT or NLS mutant GFP-Kif18B, but there was a small reduction in the average surface area of the GFP foci in cells expressing the NLS mutant relative to WT Kif18B, p<0.01 (Figure 5, B and C). In comparing the localization of Kif18B relative to EB1 in WT and NLS expressing cells, nearly 85% of the GFP foci colocalized with EB1, but only about 50% of the EB1 foci were localized with GFP (Figure 5D). Consistent with previous studies (Stout *et al*., 2011; Tanenbaum *et al*., 2011), these results confirm that Kif18B localizes with only a subset of MT plus ends. To look at the distribution of Kif18B localization, we partitioned the 3D volumes of the cells with concentric spheres and measured the distribution of EB1 and GFP-Kif18B foci in these spheres. The outer sphere was placed at the edge of the EB1 staining, and the inner sphere was 60% of the diameter of the outer sphere. We found that for both WT GFP-Kif18B and GFP-Kif18B-NLS, approximately 60% of the EB1 foci were within the inner sphere, and only about 40% of the EB1 foci were within the outer sphere (Figure 5E, left). For the GFP foci, 62% of WT GFP-Kif18B were in the outer sphere, consistent with previous observations showing that Kif18B is enriched at the ends of astral MTs. In contrast, for the GFP-Kif18B NLS, there was an even distribution of GFP foci between the outer and inner spheres where the GFP foci in the outer sphere was significantly reduced, p<0.001, suggesting that the NLS mutation hinders the proper localization of Kif18B at astral MT plus ends (Figure 5E, right). Given that the overall number of GFP foci is similar between WT and the NLS mutant, this suggests that if there is sufficient Kif18B present, it likely eventually walks to the MT plus end to regulate MT dynamics and thus aster size.

**Figure 5.**
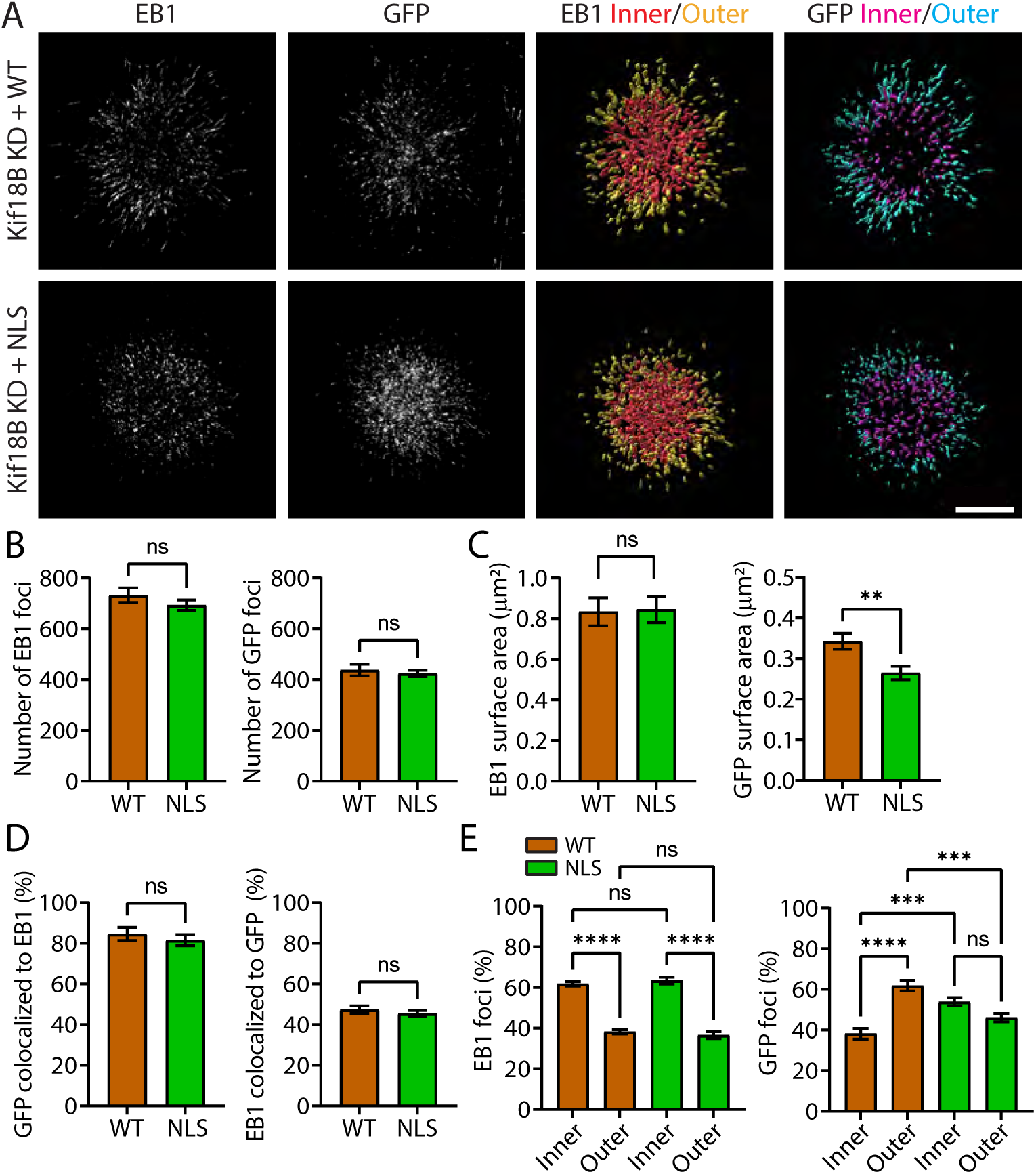
Mutation of the NLS impacts Kif18B localization. (A) Cells treated with control Neg2 or Kif18B siRNAs were mock transduced or transduced with WT or NLS mutant lentiviruses, and then fixed and stained for EB1 and GFP-Kif18B (left two panels). IMARIS software was used to identify EB1 and GFP-Kif18B foci on the asters and determine their distribution (right two panels). Scale bar, 5 µm. (B-C) The average number of EB1 or GFP foci (B) or surface areas per cell (C) is plotted as the mean ± SEM from 14 cells. An unpaired Student’s *t*-test (B, left, and C) or Mann-Whitney test (B, right) were performed to compare the means. (D) The percentage of GFP foci that are colocalized with EB1 and the number of EB1 foci that are colocalized with GFP is plotted as mean ± SEM from 14 cells. An unpaired Student’s *t*-test was performed to compare the means. (E) Two concentric rings were placed on each aster in which an outer ring was placed at the edge of the EB1 foci and a second inner ring 60% of the diameter of the outer ring was centered inside. The percentage of EB1 (3^rd^ column in A) or GFP foci (4^th^ column in A) was determined in the inner and outer spheres (color-coded in A) per cell and plotted as the mean ± SEM from 14 cells. An ordinary one-way ANOVA with the Tukey multiple comparisons test was performed to compare the means. ns, not significant; *, p<0.05; **, p<0.01; ***, p<0.001; ****, p<0.0001.

When examining the mitotic figures in Monastrol-treated cells after knockdown/rescue of Kif18B, we noticed a dramatic increase in the percentage of bipolar spindles in cells expressing high levels of WT GFP-Kif18B (Figure 6). Most control cells or cells in which Kif18B was knocked down and mock rescued had very few bipolar spindles (<5%), as would be expected for Monastrol-treated cells (Figure 6, A and B). In contrast, Kif18B knockdown cells overexpressing WT GFP-Kif18B and treated with Monastrol had ∼50% bipolar spindles (Figure 6B). These results suggest that increased MT dynamics by overexpressed Kif18B on the astral MTs promoted spindle bipolarization, which is consistent with previous studies showing that Kif18B is needed for bipolar spindle assembly in Eg5 independent cells (van Heesbeen *et al*., 2017). This role of Kif18B was proposed to be due to Kif18B regulating astral MT dynamics near the cortex and impacting cortical pulling forces that impact centrosome separation. Unlike in WT rescued cells, we did not observe dramatic increases in the bipolar spindles in cells overexpressing NLS or EBBD mutants (Figure 6B). The inability of the mutant Kif18B proteins to induce spindle bipolarization was not due to variation in expression level, as the expression of WT GFP-Kif18B and the two mutants were similar in the Kif18B knockdown cells (Figure 6C). These results suggest that while mutation of the NLS does not disrupt the ability of Kif18B to control the length of MTs in monopolar spindles, it does impair the ability of overexpressed Kif18B to induce bipolar spindle assembly in Monastrol-treated cells. This suggests that proper spatial distribution of Kif18B is important at the cortex of cells with bipolar spindles to modulate astral MT dynamics.

**Figure 6.**
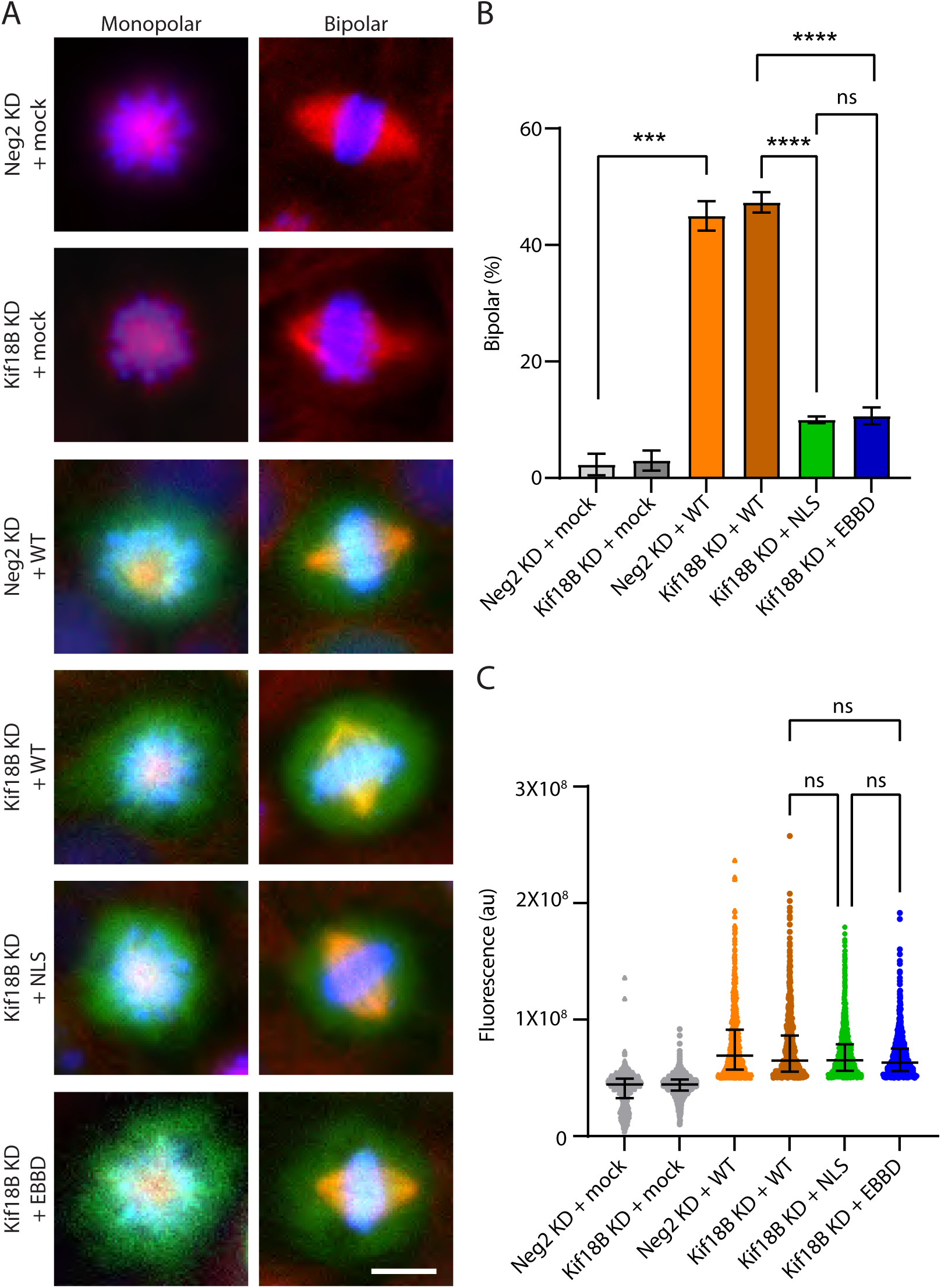
Mutation of the NLS or the EBBD inhibits bipolar spindle formation. (A) Cells treated with control (Neg2) or Kif18B siRNAs were mock transduced or transduced with WT or mutant lentiviruses, and then fixed and stained for MTs (red) and DNA (blue). Scale bar, 10 µm. (B) The percentage of bipolar spindles were determined by scoring 100 GFP-expressing mitotic cells in three independent experiments with the mean ± SEM indicated. (C) The total GFP fluorescence within 6 µm of the primary mask was calculated with an automated pipeline from three independent experiments. A minimum intensity of 5X10^7^ au was used to define GFP-Kif18B expressing cells. Each dot represents a single cell, and the median ± interquartile range is indicated. ns, not significant; ***, p<0.001; ****, p<0.0001.

### Importin α/β Increases Kif18B Association with MTs to Enhance MT Destabilization Activity

To understand how the interaction of Kif18B with importin α/β and EB1 affects the ability of Kif18B to localize to MTs and alter MT dynamics, we expressed and purified full-length WT GFP-Kif18B, added it to *in vitro* polarity marked dynamic MTs in the absence and presence of excess importin α/β or EB1, and analyzed both the motor binding events and MT lengths using a custom-built MATLAB algorithm (Figures 7A and S3A, Table S1). Previous studies showed that the plus ends of dynamic MTs assemble tubulin approximately three-folder faster than the minus end (Mitchison and Kirschner, 1984). Thus, when stabilized MT seeds are added to a solution of tubulin-GTP, MTs assemble from both ends of the seed to different lengths, resulting in the long plus-end extensions having on average three-fold longer MT extensions than the short minus-end extensions (Kristofferson *et al*., 1986). We visualized this difference in polarity by polymerizing the seeds with Cy5-tubulin (gray) and the MT extensions with X-rhodamine tubulin (magenta) (Figure 7A). Assembling polarity-marked MTs resulted in four classes of MTs: seed only, seed with one extension (1-sided), seed with two extensions of different lengths (2-sided), and extension only (Figure S3A). We assigned the long extension of 2-sided MTs as the plus end and the short extension as the minus end, and likewise we assigned the single extension of 1-sided MTs as the plus end. Of the MTs with seeds, we found approximately 50% 2-sided MTs, 35% 1-sided MTs, and 15% seed only in control and experimental conditions (Figure S3B). These ratios did not change significantly upon addition of Kif18B (Figure S3B), suggesting that subsequent statistical analysis remains robust.

**Figure 7.**
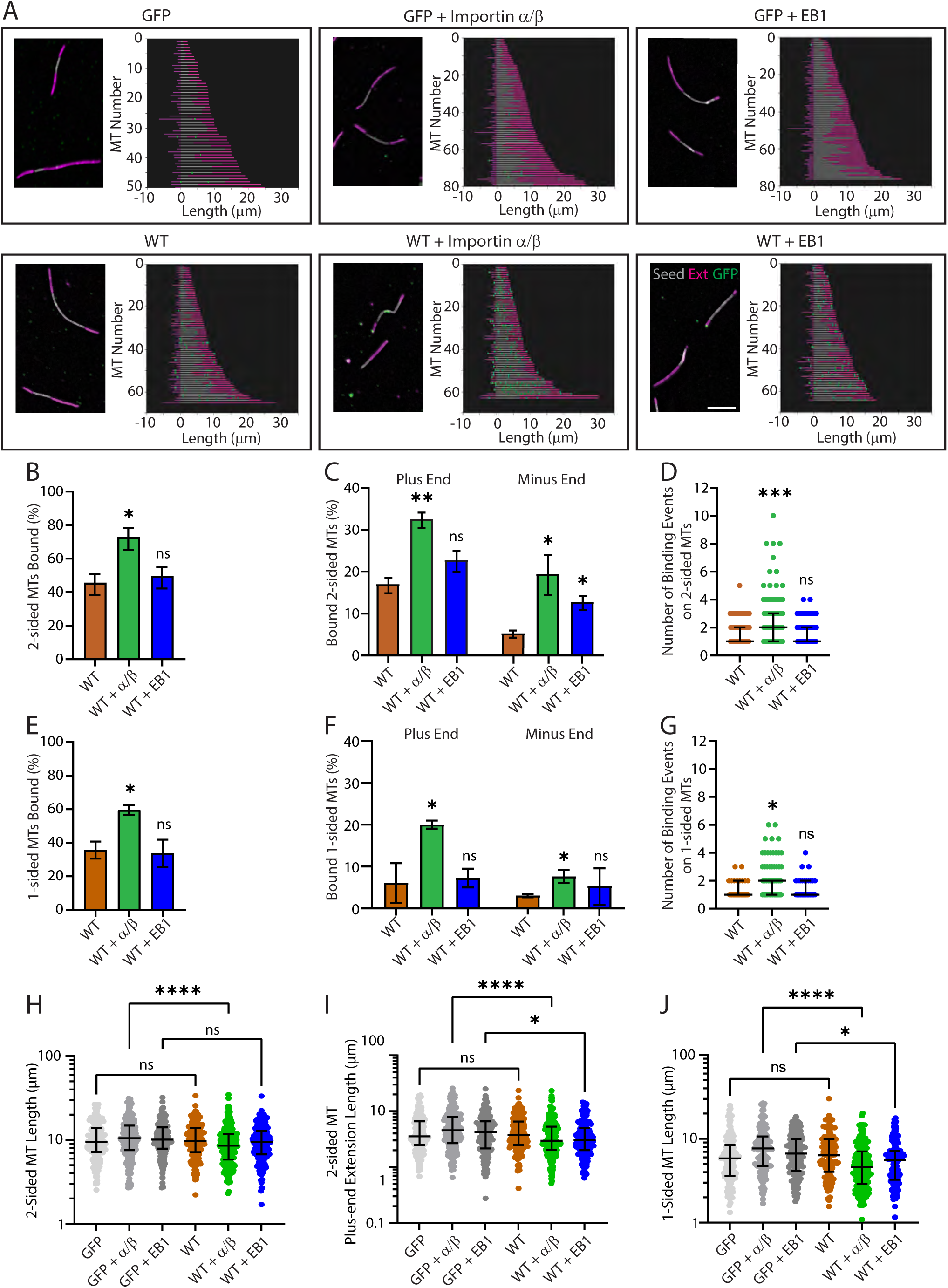
Importin α/β increase Kif18B MT association and stimulate its MT destabilization activity *in vitro*. (A) X-rhodamine labeled dynamic MTs (magenta) were nucleated from GMPCPP stabilized Cy5 MT seeds (gray) in the presence of 75 nM GFP or GFP-Kif18B ± 300 nM importin α/β or EB1, sedimented onto coverslips, and stained with anti-GFP antibody (green). GFP-Kif18B motor binding events and MT lengths were analyzed using a custom-built MATLAB algorithm. Representative 2-sided MT images (left) and the 2-sided MTs analyzed with the algorithm (right) from one of three experiments. Scale bar, 10 µm. (B and E) The percentage of 2-sided (B) or 1-sided (E) MTs with GFP-Kif18B binding were determined from each experiment and graphed as the mean ± SEM. An unpaired Student’s *t*-test was performed to compare the means. (C and F) The percentage of 2-sided (C) or 1-sided (F) bound MTs with GFP-Kif18B binding events on the distal plus or minus ends were graphed from each experiment as the mean ± SEM. An unpaired multiple *t*-test was performed to compare the means. (D, G) Of the 2-sided (D) or 1-sided (G) MTs with GFP-Kif18B motor binding events, the number of events per MT was determined and graphed from all three experiments with the indicated median ± interquartile range (n=35-174). An ANOVA with Kruskal-Wallis and Dunn’s multiple comparisons test was performed to compare the medians. (H-J) The entire lengths of the 2-sided MTs (H), the 2-sided MT plus-end extensions only (I), and the entire 1-sided MT lengths (J) were plotted for each MT on a log scale to better represent the lognormal distribution with the median ± interquartile range (n=102-248). A Kruskal-Wallis ANOVA with Dunn’s multiple comparisons test was performed to compare the medians. The number of MTs scored for MT binding and length per condition was 102-248. ns, not significant; *, p<0.05; **, p<0.01; ***, p<0.001; ****, p<0.0001.

We found that WT GFP-Kif18B localized to 46% of the 2-sided MTs, and of those MTs with motor binding events, Kif18B enriched at the distal ends of 17% of the MT plus-end extensions and only 5% of the minus-end extensions, as would be expected for a plus-end directed motor (Figure 7, B-C; Table S2). A four-fold excess addition of importin α/β to the reaction resulted in increased GFP-Kif18B binding events to 73% of the MTs, p<0.05 (Figure 7B), suggesting that the binding of importins to Kif18B may serve as a delivery mechanism for targeting Kif18B to MTs. Furthermore, the presence of importin α/β increased the percentage of binding events at the distal MT plus ends to 33% of the motor bound MTs, p<0.01 (Figure 7C). In addition to a higher percentage of MTs with motor binding events, the importins also increased the median number of binding events per MT compared to without the importins, p<0.001 (Figure 7D). In contrast, addition of EB1 did not increase the mean percentage of MTs with binding events, the mean percentage of bound MTs with distal plus-end MT binding, nor the median number of binding events per MT compared to GFP-Kif18B alone (Figure 7, B-D). Examination of the minus ends of bound MTs revealed increases in distal minus-end MT localization compared to GFP-Kif18B alone, wherein addition of the importins increased localization to 19%, p<0.05, and EB1 increased localization to 13%, p<0.05 (Figure 7C). Similar to what we found with the 2-sided MTs, importin α/β addition, but not EB1 addition, increased the percentage of 1-sided MTs with bound GFP-Kif18B, p<0.05, increased the percentage of bound MTs with plus- and minus-end MT localization, p<0.05, and increased the number of binding events per MT, p<0.05 (Figure 7, E-G). Together, these results suggest that the presence of the importins increases the association of Kif18B to MTs, which results in a net increase in plus-end MT localization.

Polymerization of dynamic MTs is stochastic and results in a non-Gaussian or lognormal distribution of MT lengths (Figures 7A and S3A). To understand how importin α/β or EB1 affect the MT destabilization activity of Kif18B, we plotted the lengths of the entire 2-sided and 1-sided MTs as well as the lengths of their seeds, plus-end extensions, and minus-end extensions on a log scale to better represent their distributions (Figures 7H-J and S3C-F). GFP-Kif18B alone did not cause a reduction in the median length of the 2-sided or 1-sided MTs, the seeds, plus-end extensions, or minus-end extensions compared to the GFP control (Figures 7H-J and S3C-F; Table S3). In contrast, addition of importin α/β stimulated GFP-Kif18B activity and resulted in a 2 μm or 19% reduction in the median 2-sided MT length, p<0.0001, and a 1.5 μm or 35% reduction in the plus-end extension length, p<0.0001, without a change in the seed or minus-end extension lengths (Figures 7, H and I and S3, C and D). Consistent with the analysis of the 2-sided MTs, GFP-Kif18B alone did not reduce the median lengths of 1-sided MTs, its seeds, or the plus-end extension (Figures 7J and S3, E and F). In addition, importin α/β stimulated GFP-Kif18B MT destabilization activity on 1-sided MTs resulting in a 3.1 μm or 40% reduction in median MT length, p<0.0001, and a 0.90 μm or 33% reduction in plus-end extension length, p<0.0001 (Figures 7J and S3F). Addition of EB1 to GFP-Kif18B did not decrease the 2-sided MT length compared to the GFP + EB1 control but resulted in a moderate decrease in plus-end extension length by 1.1 μm or 26%, p<0.05 (Figure 7, H and I), without affecting the seed or minus-end extension lengths (Figure S3, C and D). In contrast, addition of EB1 reduced the 1-sided MT length by 1.1 μm or 14%, p<0.05 (Figure 7J). The observation that EB1 did not significantly increase the percentage of MTs with motor binding events nor the percentage of distal plus-end MT binding, yet still reduced MT length, is consistent with the idea that the importins enhance the MT destabilization activity of Kif18B by a mechanism distinct from EB1.

### Importin α/β Increases Kif18B MT On-rate and Dwell Time to Increase MT Association

To understand how importin α/β affect the motile properties of Kif18B, we imaged single molecules of purified GFP-Kif18B in the absence and presence of a four-fold molar excess of importin α/β on fluorescently labeled MTs by TIRF microscopy (Figure 8A, Video 1, Table S4). Overall, we found that approximately 60% of the events were diffusive, and 40% were directed motility events (Figure S4A), consistent with one previous study but distinct from another (Shin *et al*., 2015; McHugh *et al*., 2018). The on-rate for GFP-Kif18B was 0.26 µm·s^-1^, which was increased by 36% to 0.35 µm·s^-1^ in the presence of importin α/β, p<0.05 (Figure 8B, Table S5). Once bound to the MT, Kif18B moved at a velocity of 0.26 µm s^-1^, which was not changed in the presence of importin α/β (0.25 µm s^-1^), p=0.26 (Figure 8C). The run-lengths of the translocating motors were also the same in the presence (1.18 µm) and absence (1.19 µm) of importin α/β, p=0.92 (Figure S4B). We next assessed the dwell time of the diffusive motors on the MT lattice and at the MT plus ends. GFP-Kif18B dwelled on the MT lattice for 1.77 s (corrected for photobleaching) in the absence of importin α/β, which increased by 26% to 2.30 s in the presence of importin α/β, p<0.001 (Figures 8D and S4, C and D). In contrast, there was no difference in the dwell time of GFP-Kif18B at MT plus ends without (5.56 s) or with (5.39 s) importin α/β, p=0.91 (Figure 8E). Together these data show that the addition of importin α/β promotes association of Kif18B with the MT lattice, which could then help promote MT destabilization activity when the Kif18B translocated to the MT end.

**Figure 8.**
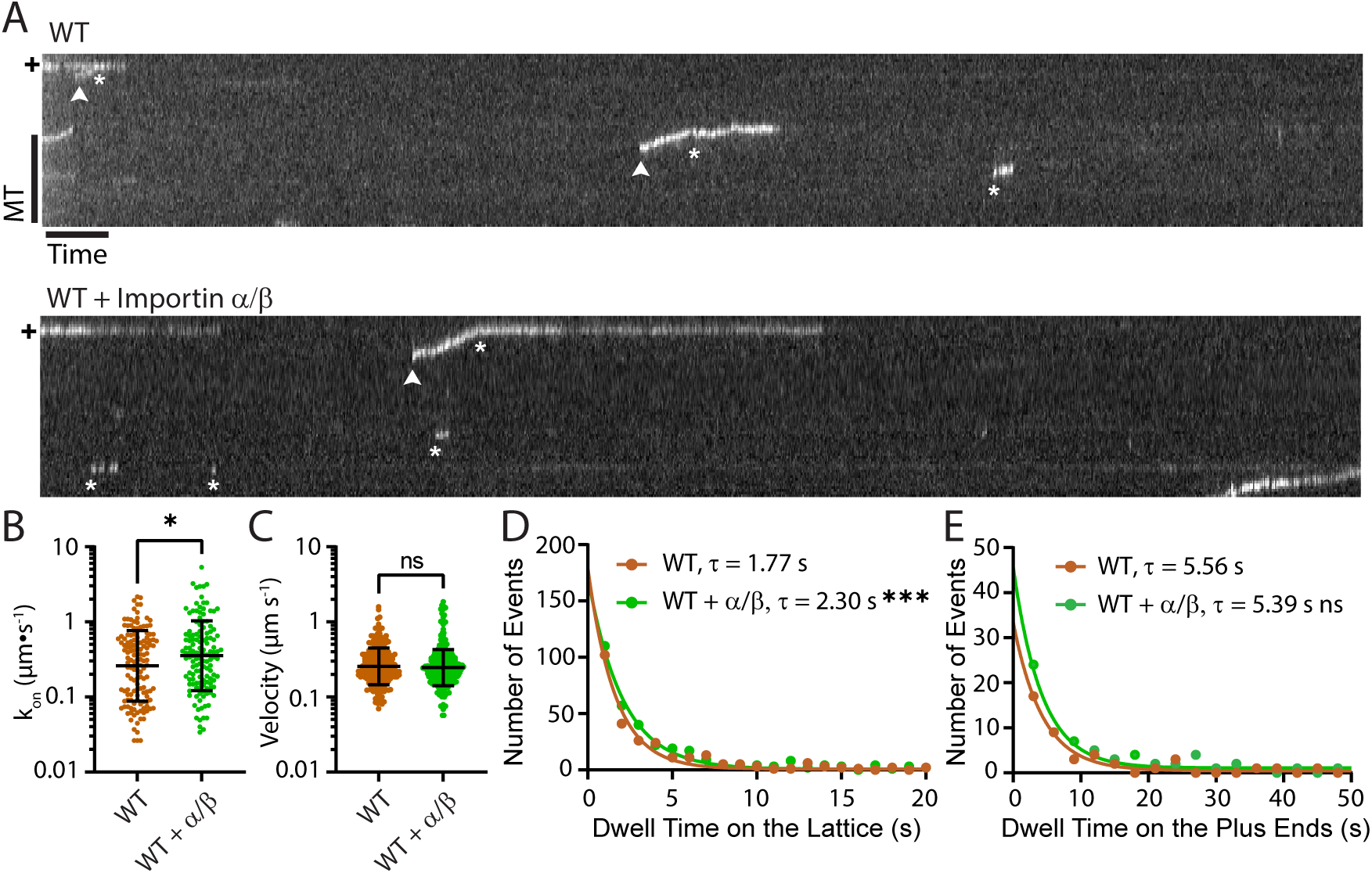
Importin α/β increase Kif18B on-rate and dwell time on the MT lattice. (A) Representative kymographs of WT GFP-Kif18B motility (top) and WT + importin α/β (bottom). The length of the MT is vertical with respect to the kymograph with the position of the plus end indicated by +, and time is horizontal. Directed moving events are indicated with an arrowhead and diffusive events or phases are indicated by an asterisk. MT scale bar, 5 µm and time scale bar, 5 s. (B and C) The on-rates (n=133-139) of GFP-Kif18B ± importin α/β binding events for each MT, *k*_on_, (B) and the velocities (n=229-346) of moving GFP-Kif18B ± importin α/β (C) are graphed from four independent experiments with the geometric mean ± the geometric SD factor indicated. A Mann-Whitney U test was performed to compare the ranks. (D and E) The dwell times of GFP-Kif18B ± importin α/β on the lattice, n=278-336, (D) or MT plus end, n=47-70, (E) are graphed as a histogram and fit with a single exponential decay model to determine the off-rates of the motor from the MT lattice or plus end. The observed off-rates were corrected for photobleaching using *k_B_*=0.0517 s^-1^, and the mean dwell time, τ, are shown and determined from the inverse of the corrected off-rate. The off-rates of GFP-Kif18B ± importin α/β were compared by an extra sum-of-squares F test. ns, not significant; *, p<0.05; ***, p<0.001.

There was an apparent contradiction in our data, as the addition of importin α/β inhibited MT binding of the Kif18B tail, whereas it promoted association of the full-length protein. One possibility is that full-length Kif18B is auto-inhibited by the tail domain akin to Kinesin-1 steric and catalytic autoinhibition (Friedman and Vale, 1999), and that importin α/β binding to the tail of full-length Kif18B protein releases this autoinhibition and promotes association of the motor with MTs. A prediction of that model is that importin α/β could then be viewed as a cargo of Kif18B and would be transported along the MT. To test this idea, we did two-color TIRF imaging of GFP-Kif18B in the presence of importin α-mCherry and unlabeled importin β. We saw no evidence of importin α-mCherry association with GFP-Kif18B either during landing events or during translocation along the MT (Figure S4E, Video 2). As a complementary approach to test the autoinhibition model, we looked at MT pelleting of GFP-Kif18B in different nucleotide states in the absence and presence of importin α/β. If Kif18B is sterically and catalytically autoinhibited by the tail, then we would expect that Kif18B should not fully bind to MTs in the absence of the importins but should bind better in their presence (Hammond *et al*., 2010). In the presence of either ATP or AMPPNP, which represent the tight binding state of the Kinesin-8 motor domain (Hunter *et al*., 2022), the majority of Kif18B pelleted with MTs (>90%), which was reduced slightly to approximately 80% in the presence of importin α/β, p<0.05 (Figure S4F). In the presence of ADP+P_i_, which is the weak binding state of the motor domain, 79% of Kif18B pelleted with MTs, which was significantly reduced to approximately 29% in the presence of importin α/β, p<0.0001. This data suggests that the majority of Kif18B association with the MT occurs through the motor domain and that importin α/β may promote this binding geometry by preventing association of Kif18B tails with MTs during landing events.

## Discussion

In this study we defined the critical regions of interaction between the Kif18B tail and its binding partners, the nuclear transport receptors importin α/β and EB1 and show how these interactions control Kif18B activity *in vitro* and in cells. We show that the importins bind directly to the tail of Kif18B and stimulate Kif18B MT destabilization activity *in vitro* by increasing the association of Kif18B with the MT without affecting its velocity. In cells, we propose that this interaction is important for delivering Kif18B to the MTs in spindle regions outside the RanGTP gradient, which facilitates Kif18B localization to the MT plus tips near the cortex. Our findings also suggest that the importins and EB1 regulate Kif18B MT destabilization in cells through distinct mechanisms, wherein the importins function to increase Kif18B MT binding and facilitate localization to the MT plus end, and EB1 functions specifically at MT plus ends for Kif18B accumulation. Overall, our findings suggest that the nuclear transport machinery (Ran and importins) is important to modulate spindle assembly factors in the vicinity of the chromosomes as well as distant from the chromatin, which may be a more general mechanism for modulating both the affinity and spatial distribution of MT binding proteins.

### EB1 and Importins Cooperate to Modulate Kif18B Activity

It is well established that the Kif18B tail binds to EB1 and that this interaction is needed for MT plus-end accumulation in cells and *in vitro* (Stout *et al*., 2011; Tanenbaum *et al*., 2011; Shin *et al*., 2015; McHugh *et al*., 2018; Dema *et al*., 2022; McHugh and Welburn, 2023), but how other regulatory factors govern the interaction of Kif18B with MTs is less clear. GFP-Kif18B is a plus-end directed motor that is mostly diffusive with short runs of directed motility that is modulated by a second MT binding domain in the tail (Shin *et al*., 2015). The presence of the tail decreases the MT off-rate, increases processivity of the motor, and increases the dwell time at the MT plus end, indicating the importance of the tail in Kif18B MT destabilization (Shin *et al*., 2015; McHugh *et al*., 2018; McHugh and Welburn, 2023). Our current work shows that blocking the interaction of Kif18B with EB1 using EB1 site-specific mutations prevented MT plus-end accumulation and MT length control in Kif18B knockdown cells, but these mutations had no effect on MT binding of the tail domain *in vitro*, indicating that EB1-mediated plus-end MT binding in cells occurs strictly through EB1-Kif18B interaction. *In vitro,* EB1 addition did not affect tail MT affinity, MT association of the full-length Kif18B protein to dynamic MTs, nor Kif18B plus-end MT localization; yet we did see a reduction in plus-end extension length in our dynamic MT seed assay, suggesting EB1 is promoting Kif18B activity specifically at MT plus ends.

Surprisingly, we saw an apparent increase in distal MT minus-end localization of Kif18B in the presence of EB1 and importin α/β, which might suggest that Kif18B also functions at MT minus ends. Consistent with this idea, a truncated Kif19A protein containing only the motor domain, neck-linker, and neck coiled-coil depolymerized MTs from the minus ends, albeit 18-fold less efficiently than from the plus end (Niwa *et al*., 2012). However, we did not see a decrease in the minus-end extension length in the presence of EB1 or importin α/β. Another possibility for the increase in MT minus-end binding events could be that Kif18B destabilized the plus-end extension to a length shorter than the minus-end extension, which then scored as a minus-end extension in our analysis. To assess this possibility, we re-categorized the distal MT minus-end binding events of GFP-Kif18B ± EB1 or importin α/β as distal MT plus-end binding events if the originally categorized plus-end extension length was less than 2.7 μm, which accounts for 90% of the control minus-end extension lengths (Figure S3G, Table S6). In the reactions with GFP-Kif18B and EB1, this re-categorization increased the number of plus-end MT binding events compared to GFP-Kif18B alone and reduced the minus-end MT binding events, suggesting that EB1 promotes Kif18B plus-end localization that in turn enhances Kif18B MT destabilization activity. Similarly, for GFP-Kif18B with importin α/β addition, this re-categorization increased the percentage of MTs with distal plus-end localization reduced the distal MT minus-end localization compared to GFP-Kif18B alone. Together, these observations support the idea that EB1 directly enhances Kif18B MT plus-end localization where it can act to regulate MT dynamics.

In cells, mutation of the importin α binding sites in the NLS mutant disrupted the distribution of Kif18B plus-end MT localization. One possibility is that the NLS mutations in the tail domain could have reduced the run length and/or dwell time of the full-length motor, resulting in reduced MT plus-end localization because the NLS mutations reduce tail-mediated MT binding. However, we do not favor this idea because the NLS and WT GFP foci number and their colocalization with EB1 were not affected, consistent with the NLS mutations having only a minor effect on the MT affinity of the tail. Alternatively, the NLS mutations could have reduced the EB1 dependent MT plus-end affinity or dwell time because we saw a reduction in the mean GFP foci surface area of the NLS mutant relative to WT GFP-Kif18B. Rather, we postulate that because importin α/β had a more pronounced effect in our dynamic MT seed assay by increasing the binding of Kif18B to MTs that resulted in higher plus-end MT localization and reduced plus-end extension length, these results indicate the importins favor motor association with MTs. Consistent with this idea, our single motor assays showed that the importins increased the on-rate and increased the dwell time on the MT lattice, which would result in a net increase in the amount of motor on the MT and ultimately the motor flux to MT ends where it can induce MT destabilization. These findings also agree with previous computer simulations of MT destabilizers on dynamic MTs, which showed that increasing the on-rate of a plus-end directed MT capping protein results in more motor binding at MT plus ends and reduced MT length (Weaver *et al*., 2011).

An apparent conundrum with our work is that the importins inhibit binding of the Kif18B tail to MTs, but they increase the overall amount of Kif18B on the MT lattice. One idea is that the Kif18B tail binds to the motor and auto-inhibits the motor from binding MTs by impeding its catalytic activity, akin to other kinesins that are autoinhibited in solution (Friedman and Vale, 1999; Imanishi *et al*., 2006; Espeut *et al*., 2008; Verhey and Hammond, 2009; Hammond *et al*., 2010). We do not favor this model, as our MT pelleting assays showed that Kif18B binds well to MTs in the absence of importins and that the importins had only a slight effect on binding of Kif18B to MTs in the presence of ATP or AMPPNP, suggesting that most of the protein binds to MTs via the motor domain rather than the tail domain. Alternatively, the importins could act as a co-factor for MT association of the motor domain, which would be consistent with how MAP7 acts to help load Kinesin-1 to MTs to modulate its activity (Ferro *et al*., 2022). However, unlike MAP7, the importins do not bind directly to the MTs, and we found no evidence of association of the importins with Kif18B while bound to the MTs.

Using AlphaFold structural modeling (Jumper *et al*., 2021), we found that the tail domain of Kif18B is largely unstructured, and that the overall molecule is more compact rather than extended like the canonical kinesin structure of motor domain followed by a long stalk and then a tail domain. Structural modeling of Kif18A and Kip3 also predicted an unstructured C-terminal tail domain. It has been proposed that the disordered tail of Kif18B may mediate interaction with partners, such as MCAK, through interaction of disordered domains (McHugh and Welburn, 2023). However, we favor the idea that importin binding to the tail of Kif18B helps a structural transition from a disordered to an ordered state of the full-length protein, which may help orient the motor domain to bind MTs and position the tail domain for its diffusive interaction with MTs (Figure 9A). The unstructured tail may reduce association of the motor domain with the MTs by steric hindrance, or the unstructured tail may bind MTs less efficiently. Binding of the importins would then cause a structural transition in the tail that now facilitates motor binding. In this model, importin binding to the tail helps promote Kif18B binding to the MT with its motor domain rather than its tail domain because the tail MT binding site is blocked by the importins. Once the motor domain binds, the importins are released, and the tail would now be free to bind MTs and act as the tether that is necessary for Kif18B diffusive or processive motility (Figure 9A). This idea is consistent with our observation that while the importins promote MT association, they are not bound to Kif18B when it is bound to the MTs. In addition, tailless versions of Kif18B are faster but less processive, and have decreased dwell times at MT plus ends (Shin *et al*., 2015). If the importins remained bound to Kif18B while moving on the MT, we would expect that in the presence of the importins that Kif18B motility would be more like that of the tailless Kif18B, which it is not. This suggests that the importins are critical for regulating association of the motor with MTs rather than regulating motor velocity or hampering plus-end MT interaction.

**Figure 9.**
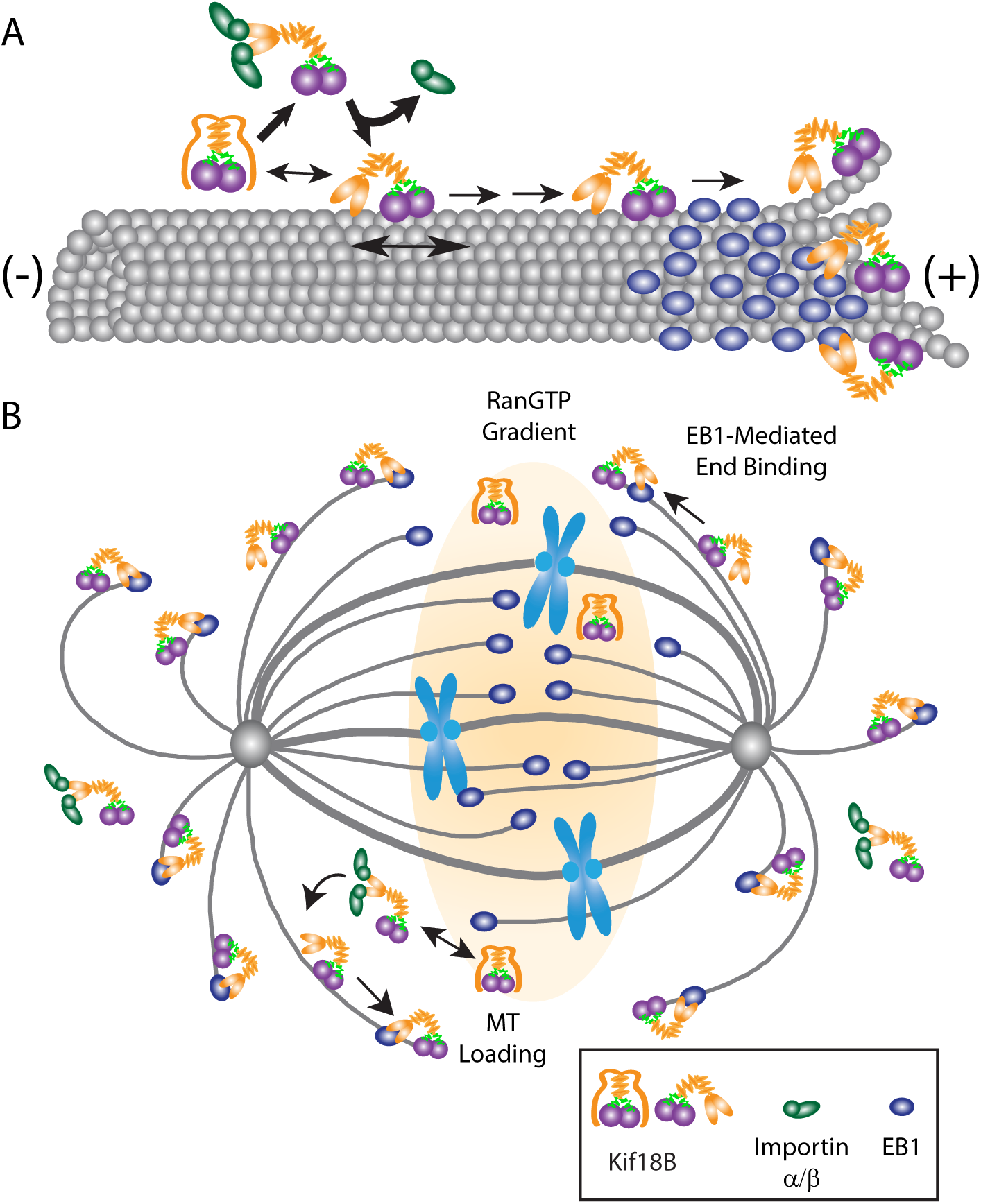
Importin α/β and EB1 differentially regulate Kif18B localization and MT destabilization activity. (A) We propose the tail of Kif18B is disordered and may sterically inhibit the motor binding to MTs. Importin α/β binding to the tail helps structure the tail and facilitates the Kif18B motor domain binding to MTs (heavy arrows). Kif18B can bind MTs independently of importin α/β but with reduced efficacy (small double arrow). The Kif18B motor can diffuse on the lattice (double arrow below the motor) or walk on the MT (directed arrows), which is modulated by the diffusive tail domain. Upon encountering a comet of EB1 on the MT plus end, the tail domain can associate with EB1 to accumulate Kif18B at the MT plus tips where it can be positioned for MT destabilization. (B) We propose that near the chromosomes, high RanGTP prevents importin α/β interaction with the tail of Kif18B, limiting Kif18B MT binding. Away from the chromatin where RanGTP levels are low, importin α/β can interact with the tail of Kif18B promoting Kif18B MT association and translocation to the plus ends (MT Loading). At MT plus ends where EB1 accumulates, the Kif18B tail can associate with EB1 (EB1-mediated End Binding), allowing Kif18B to accumulate for the modulation of MT dynamics.

### The RanGTP Gradient Spatially Controls Kif18B Localization and Activity

The spatial regulation of Kif18B is critical for its proper functioning in cells. We postulate that Kif18B is loaded onto MTs in regions of the spindle outside the RanGTP gradient through its tail interaction with importin α/β, which facilitates Kif18B association with MTs and ultimate accumulation at MT plus ends (Figure 9B). One possibility is that around the chromosomes where RanGTP is high, the Kif18B stalk and tail are disordered and hinder association of the motor with MTs. In regions farther outside the RanGTP gradient in the vicinity of the astral MTs and cell cortex, importin binding could promote motor domain MT binding. In addition, Kif18B localization is regulated by phosphorylation (Tanenbaum *et al*., 2011; McHugh *et al*., 2018), and S to A mutations in the tail domain of Kif18B changed the distribution of Kif18B so it was more enriched near chromatin (McHugh *et al*., 2018). This raises the possibility that phosphorylation in the tail domain might influence the interaction of the Kif18B tail with the importins, the MTs and/or with EB1. Our biochemical data shows that the importins compete with EB1 for binding to the Kif18B tail, but only at very high concentrations can EB1 compete with the importins. This suggests that at the MT plus ends in cells where EB1 is enriched, Kif18B interactions with EB1 could be favored, facilitating Kif18B accumulation at MT plus ends, which in turn regulates MT dynamics (Stout *et al*., 2011, this study; McHugh *et al*., 2018).

One important outcome of our studies is that the Kif18B tail interaction with the importins is likely critical for the spatial regulation of motor function. In monopolar spindles, while the NLS mutant localization is more uniformly distributed on the MTs, sufficient Kif18B reaches the ends to destabilize the MTs and control aster size, perhaps because EB1 interactions are not impeded as indicated by our *in vitro* biochemical findings. While the NLS mutant is indistinguishable from wild type in control of MT length in monopolar spindles, it is defective in inducing spindle bipolarization when overexpressed, which supports previous data showing that Kif18B is needed for bipolar spindle assembly in Eg5 independent cells (van Heesbeen *et al*., 2017). This NLS bipolarization defect suggests that either the ability of Kif18B to properly get to and enrich on MT ends of cortical MTs through increased loading onto the MT lattice through importin α/β interaction or that the spatial control of Kif18B near the cortex are needed to set up pulling forces on astral MTs that are important in spindle bipolarity.

The RanGTP system is generally utilized to govern nuclear/cytoplasmic transport. In mitosis, Ran plays a key role in the spatial control of spindle assembly factors around chromosomes (Kalab and Heald, 2008) and regulates factors at kinetochores and spindle poles. Previous findings suggest that RanGTP can control protein localization far away from the RanGTP gradient outside of the spindle. For example, NuMA-LGN localization is highest at the apical cortex of mitotic cells far from the pool of RanGTP around chromatin wherein LGN localization is reduced when chromosomes come within 2 µm of the cortex (Kiyomitsu and Cheeseman, 2012), suggesting extreme sensitivity of the NuMA-LGN complex to high RanGTP levels. While we have not observed Kif18B localization to the cortex, a recent study in keratinocytes showed that Kif18B does localize to the cortex where it colocalized with NuMA (Moreci and Lechler, 2021). Loss of either Kif18B or LGN lead to defects in spindle position that are presumably linked to disruptions in MT-cortical interactions (Blumer *et al*., 2006; Kiyomitsu and Cheeseman, 2012; Zheng *et al*., 2013). Taken together, we propose that our Kif18B NLS mutant was unable to promote spindle bipolarity in Monastrol treated cells because it lacked importin-induced MT loading on astral MTs at a distance from the RanGTP gradient that are needed for proper MT dynamics to establish pulling forces through cortical interactions. Together, these studies also support the idea that RanGTP may act as a global control center to modulate spatial regulation during mitosis, not only around chromosomes but throughout the cell.

### RanGTP and Importins Control MT Binding Affinity of Motor Proteins

Another emerging theme from our studies is that importin α/β may be a major mechanism to control the affinity of MT binding by Ran-regulated spindle assembly factors, such as the molecular motors XCTK2 (Ems-McClung *et al*., 2004; Ems-McClung *et al*., 2020) and Kif18B (this study) as well as microtubule binding proteins NuMA (Wiese *et al*., 2001) and TPX2 (Gruss *et al*., 2001; Schatz *et al*., 2003a; Schatz *et al*., 2003b). For the molecular motor proteins, each of these motors has a second microtubule binding site in the tail that is used for either cross-linking or motor processivity. For the Kif18B tail, importin α/β binding was competitive with MT binding (this study), and likewise it controlled the MT binding affinity of the Kinesin-14 XCTK2 tail (Ems-McClung *et al*., 2004). One possibility is that because MT binding sites often contain basic residues similar to canonical NLS motifs (Conti *et al*., 1998; Kosugi *et al*., 2009) that their motifs coincidentally overlap. A more interesting possibility is that in proteins with an NLS, the NLS region may serve as a MT binding domain with a distinct binding site on the MT itself. Structural studies are beginning to reveal that different motors and MAPs bind to distinct regions of the MT surface (Nogales and Zhang, 2016) and that these unique binding regions may help explain how multiple MAPs can bind to a MT. This binding geometry could be important to load the motors onto the MT. Alternatively, tail MT binding may be important to position the motor head either to a different tubulin dimer on the same MT or, in the case of cross-linking motors, to another MT. Future structural studies identifying how the NLS-containing motor tails bind MTs will be important to dissect this idea.

## Materials and Methods

### Cloning and site-directed mutagenesis

To express His_6_-EB1-CyPet, full length human EB1 was amplified by PCR from pET30a-EB1 that encodes His_6_-S-EB1 (Stout *et al*., 2011), a gift from Jennifer Tirnauer) using the primers in Table S7. The resulting PCR product was cloned into the *Xba*I*/Xho*I sites of pRSETB-CyPet (Ems-McClung *et al*., 2020) to create pRSETB-EB1-CyPet. For expression and purification of the C-terminal tail proteins of human Kif18B (Kif18B) fused to YPet, we generated plasmids with an N-terminal YPet tag and a C-terminal His_6_ tag in two steps. We first generated plasmids encoding YPet fused to the full length Kif18B tail (aa 605-828) amplified from pDEST15-Kif18B(CT605-828) and the far C-terminus (aa 708-828) amplified from pDEST15-Kif18B(CT708-828) (Stout *et al*., 2011) using the primers in Table S7. The resulting PCR fragments were cloned into pRSETB-YPet (Ems-McClung *et al*., 2020) digested with *Sac*II*/Pst*I creating pRSETB-YPet-Kif18B(605-828) and pRSETB-YPet-Kif18B(708-828). To generate plasmids with a C-terminal His_6_ tag, DNAs encoding YPet-Kif18B(605-828) and YPet-Kif18B(708-828) were PCR amplified from the pRSETB-YPet constructs using primers in Table S7 and cloned into the pET24b(+) *Bam*HI*/Sal*I sites to generate pET24b(+)-YPet-Kif18B(605-828) and pET24b(+)- YPet-Kif18B(708-828), respectively. To generate pET24b(+)-YPet-Kif18B(605-707), DNA encoding YPet-Kif18B(605-707) was PCR amplified from pRSETB-YPet-Kif18B(605-828) using the primers in Table S7 and cloned into the pET24b(+) *Bam*HI*/Sal*I sites. The expression vector for His_6_-importin α-mCherry was constructed by PCR amplification of mCherry with primers listed in Table S7 and subcloned into the *Kpn*I/*Hind*III sites of His_6_-importin α-CyPet after dropout of the CyPet (Ems-McClung *et al*., 2020). To generate the plasmid for expression of His_6_-mCherry, the coding sequence of mCherry was amplified with primers listed in Table S7 and subcloned into the *BamH*I/*Sac*I sites of pRSETA.

Two putative nuclear localization signal (NLS) sequences in the tail of Kif18B were predicted using the web-based predication program cNLS Mapper (http://nls-mapper.iab.keio.ac.jp/cgi-bin/NLS_Mapper_form.cgi). The putative non-canonical bipartite sequence (NLS1) was further evaluated using guidelines described by (Kosugi *et al*., 2009) to determine the residues critical for importin α binding. The codons encoding the NLS1 basic residues, indicated in bold, ^639^**KKR**RRPSALEADSPMAP**KR**^657^ were mutated sequentially to alanine with the NLS1a and NLS1b primer sets listed in Table S8 using the QuikChange site-directed mutagenesis system (Agilent, San Diego, CA) to create pET24b(+)-YPet-Kif18B(605-828)-NLS1. Likewise for the second putative NLS (NLS2), the codons encoding the basic residues of the putative monopartite sequence ^795^**KKKR**^798^ were mutated to alanine with the primers in Table S2 to create pET24b(+)-YPet-Ki18B(605-828)-NLS1/2 and pET24b(+)-YPet-Kif18B(708-828)-NLS2, respectively. To generate pET24b(+)-YPet-Kif18B(605-707)-NLS1 containing only the NLS1 mutations, DNA was amplified from pET24b(+)-YPet-Kif18B(605-828)-NLS1 and cloned into the *Bam*HI*/Sal*I restriction sites of pET24b(+).

To identify the EB1 binding sites in the tail of Kif18B, the codons encoding the last two residues, indicated in bold, of the three previously reported EB1 binding sites (Tanenbaum *et al*., 2011), ^665^SF**LP**^668^ (E1), ^786^SS**LP**^789^ (E2), and ^812^AR**LP**^815^ (E3), were mutated to asparagine (Honnappa *et al*., 2009) using the QuikChange site-directed mutagenesis system and the primers listed in Table S8. Double EB1 mutants were generated by sequential site-directed mutagenesis. All constructs were verified by sequencing.

### Protein expression and purification

For the expression of His_6_-EB1-CyPet, His_6_-S-EB1 (Stout *et al*., 2011), His_6_-importin α-CyPet, His_6_-S-importin β, and importin α-His_6_ (Ems-McClung *et al*., 2004), His_6_-importin α-mCherry, and His_6_-mCherry recombinant proteins, plasmids were transformed into BL21(DE3)pLysS bacteria and induced with 0.1 mM isopropyl β-D-1-thiogalactopyranoside (IPTG) at 16°C or 20°C for 24 h. For the expression of YPet-Kif18B-His_6_ recombinant proteins, plasmids were transformed into SHuffle T7 Express LysY bacteria (NEB Cat# C3030J) and induced with 0.4 mM of IPTG at 16°C for 24h. Cells were pelleted, frozen in liquid nitrogen, and stored at -80°C. For protein purification using NiNTA agarose (Qiagen), cells were lysed in 50 mM phosphate buffer, pH 8.0, 300 mM NaCl, 10 mM imidazole, 0.1% Tween-20, 1 mM PMSF,1 mM benzamidine, 0.5 mg/ml lysozyme and sonicated four to six times using a Branson Sonifer at 20% output. Immediately after sonication, the salt concentration was increased to 500 mM by adding NaCl, and β-mercaptoethanol (β-ME) added to a final concentration of 5 mM. The lysate was clarified at 18,000 rpm for 20 min at 4°C in a Beckman JA25.5 rotor. The supernatant was added to NiNTA agarose equilibrated with Column Buffer (50 mM phosphate pH 8.0, 500 mM NaCl, 0.1% Tween-20, 10 mM imidazole) in a 50 ml falcon tube and incubated at 4°C for 1 h with rotation. The clarified lysate-bead mixture was centrifuged in a clinical centrifuge for 4 min at ∼400 *g* to pellet the beads, which contained the bound protein of interest. After discarding the supernatant, the agarose beads were washed twice in 10 column volumes (CV) of Column Buffer containing 1 mM PMSF, 1 mM benzamidine, 5 mM β-ME for 10 min with rotation. Ten CV of Column Wash Buffer (50 mM phosphate buffer, pH 6.0, 500 mM NaCl, 5 mM β-ME, 0.1 mM PMSF) was added to the pelleted beads and loaded into a disposable column at 4°C. Once the Column Wash Buffer was drained, the bound protein was eluted with 10 CV of Elution Buffer (50 mM phosphate buffer, pH 7.2, 500 mM NaCl, 400 mM imidazole, 5 mM β-ME) at 4°C in 1 mL aliquots. For the purification of YPet-Kif18B-His_6_ tail proteins, 1 μg/ml leupeptin, 1 μg/ml pepstatin, 1 μg/ml chymostatin, and 1 mM PMSF were added to all the purification buffers. For His_6_-EB1-CyPet, His_6_-importin α-CyPet/mCherry, His_6_-S-importin β, importin α-His_6_, and His_6_-EB1, eluted fractions containing the highest amounts of protein were dialyzed into XB dialysis buffer (10 mM HEPES, pH 7.2, 100 mM KCl, 25 mM NaCl, 50 mM sucrose, 0.1 mM EDTA, 0.1 mM EGTA) overnight at 4°C. The proteins were further dialyzed for an additional 3 hours the next morning. YPet-Kif18B-His_6_ proteins were similarly dialyzed into XB500 (10 mM HEPES, pH 7.2, 500 mM KCl, 50 mM sucrose, 0.1 mM EDTA, 0.1 mM EGTA). The dialyzed proteins were snap-frozen in liquid nitrogen in single use aliquots and stored at -80°C.

### Förster resonance energy transfer (FRET) of Kif18 interaction with importin α/β or EB1

The interactions between human Kif18B and human importin α/β or EB1 were reconstituted *in vitro* by using the FRET biosensors, His_6_-importin α-CyPet, His_6_-EB1-CyPet, and YPet-Kif18B-His_6_ tail domains in solution as described previously (Ems-McClung and Walczak, 2020). To determine saturable binding, 100 nM of His_6_-importin α-CyPet together with 400 nM His_6_-S-importin β or 100 nM of His_6_-EB1-CyPet were mixed with 100 nM-1.0 µM YPet tagged Kif18B tail fragments in Reaction Buffer (10 mM HEPES pH 7.2, 55 mM ionic strength, 50 mM sucrose, 0.1 mM EDTA, 0.1 mM EGTA, 0.1 mg/ml casein). Where indicated, RanQ69L was added to a final concentration of 10 μM to the YPet-Kif18B tail + importin α-CyPet/ His_6_-S-importin β reactions after an initial FRET measurement, incubated for 10 min at RT, and FRET measured again (Ems-McClung and Walczak, 2020). To achieve a final ionic strength of 55 mM, XB dialysis buffer was added to the reactions accounting for the KCl and NaCl contributed by the different proteins. A 50 µl aliquot of each reaction mix was transferred to a Costar #3964 96-well ½ area black plate and incubated at room temperature for 10 min before performing a spectral scan in a Synergy H1 plate reader (Bio-Tek). The spectral scan was performed by exciting His_6_-importin α-CyPet or His_6_-EB1-CyPet at 405 nm and recording the emission from 440 to 600 nm with 5 nm steps (Ems-McClung and Walczak, 2020). The recorded data was processed in Excel by subtracting the background YPet emission in control reactions without CyPet protein from YPet emission in the presence of CyPet protein, and the net emission was then normalized to the CyPet emission at 460 nm to obtain FRET ratios at 530 nm. Graphs of the FRET ratio means ± SEM were plotted relative to the different molar ratios of importin α-CyPet and hEB1-CyPet in Prism (GraphPad). For competition assays, a premix comprising 100 nM His_6_-importin α-CyPet, 400 nM of His_6_-S-importin β and 400 nM His_6_-S-EB1 or 100 nM His_6_-hEB1-CyPet, 400 nM of importin α-His_6_, and His_6_-S-importin β were prepared and then added to 800 nM of YPet-Kif18B-His_6_ tail fragments in Reaction Buffer and processed for FRET as described. For competition assays with 30x and 40x His_6_-S-EB1, 3 µM, or 4 µM His_6_-S-EB1 were mixed with 100 nM His_6_-importin α-CyPet, 400 nM of His_6_-S-importin β, and 600 nM of YPet-Kif18B(605-828)-His_6_ in Reaction Buffer and processed for FRET as described.

Full-length human Kif18B N-terminally tagged with enhanced green fluorescent protein (EGFP), GFP-Kif18B, was expressed in Sf-9 insect cells at an MOI of 2 from baculovirus generated from pFB-EGFP-Kif18B-siRNA^R^ using Gateway technology with pENTR-EGFP-Kif18B-siRNA^R^ (see Lentivirus Production below) and pFastBac1-DEST plasmids. pFastBac1-DEST was generated by amplifying the destination cassette (ccdB and chloramphenicol resistance genes) from pDEST^TM^15 (ThermoFisher) with flanking attR1/R2 sites (Table S7) and inserting them into the SacI/KpnI sites of pFastBac1. Expressed GFP-Kif18B was purified by cation exchange chromatography using a 1 mL HiTrap SP column followed by size exclusion using a Superose 6 HR 10/30 column in 20 mM Pipes pH 6.8, 300 mM KCl, 1 mM MgCl_2_, 1 mM EGTA, 0.1 mM EDTA, 1 mM DTT, 0.1 mM ATP, 0.1 µg/mL LPC. Solid sucrose was added to 10%, aliquoted, frozen in liquid nitrogen, and stored at -80°C (Ems-McClung *et al*., 2004).

### Lentivirus production

To produce lentivirus for expression of human Kif18B N-terminally tagged with EGFP, GFP-Kif18B, containing the NLS and EBBD mutations in mammalian cells, pLOVE constructs containing siRNA resistant (siRNA^R^) Kif18B were made using Gateway technology. To generate pENTR-EGFP-Kif18B-siRNA^R^, the EGFP-Kif18B sequence from pEGFP-Kif18B (Stout *et al*., 2011) was amplified using the primers in Table S7, cloned into pENTR/D-TOPO (Invitrogen), and silent mutations introduced to confer resistance to siRNA of Kif18B by QuikChange site-directed mutagenesis using the primers in Table S8. Subsequent rounds of site-directed mutagenesis were used to create pENTR-EGFP-Kif18B-siRNA^R^-NLS1/2 with the pENTR-Kif18B-NLS1a, pENTR-Kif18B-NLS1b, and pENTR-Kif18B-NLS2 forward and reverse primers and pENTR-EGFP-Kif18B-siRNA^R^-EBBD1/3 with the pENTR-Kif18B-EBBD1 and pENTR-Kif18B-EBBD3 forward and reverse primers in Table S8. Entry vectors were recombined into the pLOVE destination vector (Addgene #15948) using Gateway technology. All clones were verified by sequencing.

HEK293T cells were maintained in Gibco™ DMEM, High Glucose supplemented with GlutaMAX™ and pyruvate (Thermofisher), and 10% fetal bovine serum (Corning) at 37°C and 5% CO_2_ in a humidified incubator. Lentiviruses were produced by co-transfecting 1 μmol of pLOVE-EGFP-Kif18B-siRNA^R^, pLOVE-EGFP-Kif18B-siRNA^R^-NLS1/2, or pLOVE-EGFP-Kif18B-siRNA^R^-EBBD plasmids with 1 μmol each of the packaging plasmids dRT-pMDLg/pRRE (Addgene #60488), pRSV-Rev (Addgene #12253), and VSV-G envelope expressing plasmid, pMD2.G (Addgene #12259) into HEK293T cells plated at 1.5 x 10^5^ cells/ml in T-25 flasks using Lipofectamine 3000 (Thermo Fisher). Media was changed 6-8 h post-transfection, and media containing virus was collected at 24 h and 48 h post-transfection, filtered, and stored at -80°C in single use aliquots. Functional titers of these viruses were determined post 48h of viral transduction as described in <u>Addgene: Fluorescence</u> <u>Titering Assay</u> using 96-well clear bottom plates (Corning #3603).

### Kif18B siRNA knockdown and EGFP-Kif18B-siRNA^R^ lentiviral expression

HeLa cells were maintained in Gibco™ DMEM, High Glucose supplemented with GlutaMAX™, pyruvate, and 10% fetal bovine serum at 37°C and 5% CO_2_ in a humidified incubator. For Kif18B knockdown and/or EGFP-Kif18B-siRNA^R^ lentiviral expression assays, 5 x10^4^ HeLa cells were seeded on coverslips in 6-well plates. After 24 h, cells were transfected with 40 nM RNAi oligonucleotides using Lipofectamine RNAiMax (Invitrogen) according to the manufacturer’s instructions. The following siRNAs (Dharmacon) were used: Neg2 (5’-UGGUUUACAUGUUGUGUGA-3’) or Kif18B siRNAi (5’-CAGUUUCCAUGAAUGCAUUUU-3’) (Stout *et al*., 2011). After 24 h, the transfection media was removed, and the HeLa cells were transduced with lentivirus particles with a MOI of 0.5 in presence of 10 μg/ml of polybrene in complete media without antibiotics. To form monopolar spindles, lentivirus media was removed 36 h post infection and replaced with fresh media containing 100 µM Monastrol and incubated for 12 h prior to fixation.

### Western blot

To probe Kif18B depletion by Western blot, 120,000 HeLa cells were seeded per well in a 6 -well plate and after ∼24 h, cells were transfected with control siRNA (Neg2) or Kif18B siRNA. Cells were released from transfection after 24 h, allowed to grow for 6 h and then synchronized by treating with 6.0 µM Cdk1 inhibitor, RO3306 (Sigma, #SML0569) for 16 h followed by 100 µM Monastrol treatment for 4 h. Cells were trypsinized and resuspended in phosphate buffered saline (PBS: 137 mM NaCl, 2.7 mM KCl, 10 mM Na_2_HPO_4_, 1.8 mM KH_2_PO_4_, pH 7.2) containing 100 µM Monastrol. 50,000 and 25,000 cells in 2X sample buffer were loaded on to 10% SDS-PAGE, transferred to nitrocellulose membrane (Biotrace, #66485) for Western blot analysis. The blot was blocked in 5% non-fat dry milk in TBS-T (20 mM Tris, pH7.5, 150 mM NaCl with 0.1% Tween-20), probed with affinity purified rabbit anti-Kif18B (2.5 µg/ml) in Ab-Dil-T ((20 mM Tris, pH 7.5, 150 mM NaCl, 2% BSA, 0.1% Tween-20) as a primary antibody. Donkey anti-rabbit IgG HRP (Thermo Fisher Scientific, #45000682) (1:10,000) in 5% nonfat dry milk in TBS-T was used as a secondary antibody. Blots were incubated with SuperSignal West Pico Chemiluminescent Substrate (ThermoFisher Scientific, Rockford, IL) and exposed to Amersham Hyperfilm ELC (GE Healthcare). The film was developed using SRX-101A, Medical Film Processor (Konica Minolta Medical and Graphic, Inc).

### Immunofluorescence

For immunofluorescence, cells were rinsed in phosphate-buffered saline (12 mM phosphate, 137 mM NaCl, 3 mM KCl, pH 7.4) and then fixed in -20°C MeOH for 10 min. Cells were blocked in Abdil-Tx (20 mM Tris, 150 mM NaCl, pH 7.5 (TBS), 0.1% Triton X-100, 2% bovine serum albumin, 0.1% sodium azide) for 30 min at room temperature (RT). Cells were stained with rabbit anti-GFP (2.5 µg/ml) and mouse DM1α (1:500, Sigma-Aldrich T6199), rabbit anti-Kif18B serum (1:2000) or mouse anti-EB1 (0.5 µg/ml, BD Transduction Laboratories) primary antibodies for 30 min at RT, washed with TBS-Tx (TBS with 0.1% Triton X-100), and then stained with Alexa Fluor 488 donkey anti-rabbit (2 µg/ml, Invitrogen #A21207) and Alexa Fluor 594 goat anti-mouse (2 µg/ml, Invitrogen #A10239) secondary antibodies for 30 min at RT. DNA was stained with 10 μg/ml Hoechst for 5 min (Sigma-Aldrich) after being washed with TBS-Tx and then mounted in Prolong Diamond (Invitrogen #P36961).

For the analysis of bipolar spindles in Monastrol-treated cells (Figure 6), 4 x10^3^ HeLa cells were seeded in a 96-well clear bottom plate (Corning #3603). After 24 h, cells were transfected with 40 nM RNAi oligonucleotides using Lipofectamine RNAiMax as above. After 24 h, the transfection media was removed, and the cells were infected with lentivirus particles in complete medium containing 10 μg/ml polybrene. To form monopolar spindles, lentivirus media was removed 32 h post infection and replaced with fresh media containing 100 µM Monastrol and incubated overnight prior to fixation. Cells were fixed in -20°C MeOH with 1% formaldehyde for 5 min, blocked, and stained with mouse DM1α (1:500) for primary antibodies as described above. Alexa Fluor 594 goat anti-mouse (2 µg/ml) was used as a secondary antibody. DNA was stained with 10 µg/ml Hoechst, and then the plates were stored in PBS before imaging.

### Microscopy

For analysis of aster area, images were collected with a 40X 1.0 PlanApo objective or with a 60X 1.42 PlanApo objective for analysis of GFP localization on a Nikon 90i equipped with a CoolSnap HQ CCD camera (Photometrics) and controlled by Metamorph (Molecular Devices). Image stacks were collected through the entire cell volume at 0.4 μm and 0.2 µm steps for 40X and 60X objectives respectively. Each individual channel was imaged with equivalent exposure times across all samples. For the analysis of Kif18B localization on monopolar spindles, cells transduced with GFP-Kif18B WT or NLS mutant lentiviruses were imaged using the OMX SR 3D-SIM Super-Resolution system (GE DeltaVision) controlled by AquireSR software. Images were captured at 0.125 µm step size with an PlanApo N 60X 1.42 NA oil PSF objective, using 1.518 immersion oil. The 405 nm wavelength images were acquired at 1% laser strength, 488 nm wavelength images at 3% laser strength, 564 nm wavelength images at 4% laser strength with using exposure times that were adjusted to obtain roughly 3000 grey levels to account for differences in expression levels of GFP. Cells with equivalent exposure times were paired for analysis. Images were processed using softWoRx (Applied Precision, Inc.) and IMARIS 3D imaging software (Bitplane Inc.).

### Image analysis and quantification

Composite images were generated and processed identically using FIJI. Aster areas were measured in FIJI by manually drawing a circle around each MT aster in the Alexa 594 channel and then transferring that ROI to the 488 nm channel. To determine the lower limit of GFP fluorescence for analysis, cells with either control or Kif18B knockdown were mock-transduced and then stained with anti-GFP antibodies as described above, and the average fluorescence in the 488 nm channel was considered background for the lower GFP fluorescence threshold. To determine the upper level of the GFP fluorescence that could be used to assess the level of rescue of our WT and mutant constructs, we systematically tested bins of GFP fluorescence of cells with knockdown of Kif18B that were rescued with WT GFP-Kif18B lentivirus expression and used the broadest bin size that resulted in full rescue of the knockdown phenotype, which corresponded to a range of 40-2000 arbitrary units (au).

For analysis of the percentage of bipolar spindles and GFP expression levels in cells with overexpression of lentiviral GFP-Kif18B proteins, images were collected using a 20X 0.45 Olympus Plan Fluorite objective on a Lionheart FX automated microscope (BioTek). A semi-automated analysis pipeline was created using the Gen5 Image Prime software (BioTek) to measure the area of monopolar spindles and the intensity of the GFP signal. Briefly, primary masks for mitotic cells were defined by DAPI fluorescence above 50,000 au with a secondary mask of the 594 nm or the 488 nm channel extending up to 6 µm out from the primary mask with a threshold of 9,000 au or 10,000 au, respectively. Monopolar spindles were defined as having a length/width ratio of less than or equal to 1.25 and an area between 50-300 µm^2^. GFP-Kif18B expressing cells were defined as having a GFP intensity greater than 5 x10^7^ au. The percentage of bipolar spindles were measured by counting 100 mitotic figures and scoring the number of monopolar and bipolar spindles from three independent experiments.

Fluorescence values and aster areas were exported to Excel (Microsoft), plotted in Prism 9 (GraphPad Software), and graphs assembled in Illustrator (Adobe). Statistical analyses were performed using Prism 9. To determine if the data came from a normal distribution, a D’Agostino-Pearson omnibus normality test was performed, and p<0.05 was considered as a non-normal distribution. For comparisons of non-normally distributed GFP fluorescence and aster area, one-way analysis of variance (ANOVA) followed by a Kruskal-Wallis tests with Dunn’s multiple comparisons tests were used. Fluorescence values and aster areas were plotted with the median ± interquartile range. The mean percentages of bipolar cells were plotted ± SEM. Unpaired *t*-tests were used to determine differences between the means. A p<0.05 was considered significant for all tests.

OMX images of WT and NLS rescued cells (14 cells each), with equivalent exposure time and fluorescence values were used for the quantitative assessment of GFP and EB1 signals using IMARIS software. Briefly, GFP and EB1 objects were created using surface creation feature. The smallest diameter of the object was set to 0.25 µm for both GFP and EB1 signals (Stout et al., 2011). The images were subjected to equivalent threshold to best fit the image data. Then, the automatic threshold value for the quality, which accounts both the intensity and area of the signals, was used for both WT and NLS mutant rescued cells. Data were acquired to determine the number of foci, surface area of the foci, and the extent of colocalization of GFP with EB1 and vice versa. An unpaired Student’s *t*-test or a Mann-Whitney test were used to compare the means and p<0.05 was considered significant.

To determine the distribution of the EB1 and GFP foci, an object of 0.1μm was created at the center of the aster. The outer region of interest was defined by assigning a sphere that encompassed >98% EB1 foci. The inner region of interest had a diameter of 60% of the diameter of the outer sphere. The number of GFP and EB1 foci were obtained in the two concentric spheres. An ordinary one-way ANOVA with the Tukey multiple comparisons test was performed to compare the means, and p<0.05 was considered significant for all tests.

### Microtubule Binding and Dynamic MT Seed Assays

The interaction between Kif18B tail and MTs was measured by co-sedimentation. Doubly stabilized MTs were polymerized by incubating 5 µM tubulin with 0.5 mM GMPCPP (Jena BioScience, #NU-405L) for 20 min at 37°C, adding paclitaxel to a final concentration of 10 µM and incubating for an additional 10 min at 37°C. MTs were pelleted at 45,000 rpm in a Beckman TLA100 rotor and then resuspended in BD Buffer (80 mM K-Pipes pH 6.8, 1 mM MgCl_2_, 1 mM EGTA, 1 mM DTT) + 50 mM KCl containing 10 µM paclitaxel. The MT concentration was determined using a molar extinction coefficient of 115,000 M^-1^cm^-1^ and adjusted to 10 µM with BD + 50 mM KCl. For binding reactions, 250 nM YPet-Kif18(605-828) WT or mutant derivatives were combined with 0 to 5 µM MTs, incubated for 15 min at RT and then sedimented at 45,000 rpm in a TLA 100 rotor at 25°C. The supernatant was removed to a fresh tube, and the pellet was resuspended in an equivalent volume of BD + 50 mM KCl. A 20 µl aliquot of the corresponding supernatant and pellet fractions were transferred to a Nunc 384 well plate (#262260), and the fluorescence of YPet was measured with 500 nm excitation and 530 nm emission on a BioTek Synergy plate reader. For competition experiments, a premix solution of 250 nM YPet-Kif18(605-828) was incubated with 500 nM importin α-His_6_, and His_6_-S-importin β for 5 min before adding to the MT mixture. For competition with unlabeled EB1, the premix solution contained 250 nM YPet-Kif18B(605-828) and 1 mM His_6_-S-EB1. All data was corrected for the fraction of YPet-Kif18B tail that pelleted in the absence of MTs and normalized to represent fractional binding in micromolar protein bound. The average protein bound at each MT concentration were plotted as the mean ± SEM in Prism and fit with to the quadratic equation (Hertzer *et al*., 2006).

For MT binding of full-length GFP-Kif18B in different nucleotide states, 200 nM GFP-Kif18B ± 800 nM importin α-His_6_/His_6_-S-importin β were incubated with 1 µM doubly stabilized MTs in BD Buffer, 50 mM KCl, 5 µM paclitaxel, 0.5 µg/µl casein containing 2 mM MgATP, 5 mM MgAMPPNP, or 2 mM MgADP + 20 mM P_i_ for 20 min at RT and then sedimented at 45,000 rpm in a TLA 100 rotor at 22°C. The supernatant was removed to a fresh tube, and the pellet was resuspended in an equivalent volume of BD, 50 mM KCl, 5 µM paclitaxel, 0.5 µg/µl casein. A 15 µl aliquot of the supernatants and pellets were transferred to a Nunc 384 well plate, and the GFP fluorescence measured with 470 nm excitation and 510 nm emission on a Synergy H1 plate reader. The fraction bound was determined in Excel and plotted in Prism as the mean ± SEM from three to five independent experiments. Student’s *t*-tests were performed in Excel and significance was considered for p<0.05.

Dynamic MT seed assays were performed by polymerizing GMPCPP stabilized Cy5-MT seeds from 4 µM HiLyte 647 tubulin (Cytoskeleton, #TL670M) in BD buffer + 0.5 mM GMPCPP at 37°C for 1 hour. Seeds were added 1:10 for final concentrations of 75 nM GFP protein ± 300 nM importin α/β or 300 nM EB1, 6 µM 17% X-rhodamine tubulin, 50 mM KCl, 2 mM ATP, 1 mM GTP in BD buffer, and incubated at 37°C for 20 min. Reactions were fixed with 40 µL of 1% glutaraldehyde/BD buffer for 3 min and diluted with 800 µL BRB80 (BD buffer without 1 mM DTT). Fixed and diluted samples were added to spin down tubes containing a removable chock, coverslip, and 5 mL BRB80 underlaid with 10% glycerol, BRB80. The spin down tubes were centrifuged for 45 min at 12,000 rpm in a JS13.1 Beckman rotor at 20°C. The coverslips were removed, immunostained with 3 µg/mL anti-GFP antibody and 2 µg/mL Alexa488 donkey anti-rabbit IgG, mounted in ProLong^TM^ Diamond, cured for 24 h, and sealed with nail polish. Widefield FITC, TRITC, and Cy5 fluorescence images were taken on a Nikon A1 microscope equipped with a PlanApo VC 60x NA1.4 DIC NS objective and ORCA-Flash4.0 Hamamatsu camera controlled by Nikon Elements in the Light Microscopy Imaging Center at IU – Bloomington. Each image was taken at identical exposure times for each of the three channels, scaled identically in FIJI, and assembled in Adobe Illustrator.

A graphical user interface (GUI) was developed in MATLAB for interactive identification of MT seeds, extensions, and motors from three images taken with different fluorescence filter sets. MT polymer segmentation and identification was accomplished using a Hessian ridge algorithm (Steger, 1998) to identify contiguous fluorescent structures followed by a Dijkstra minimum path algorithm (Dimas, 2022) for tracing continuous points that minimize curvature and gaps. Parameters for Gaussian filters, maximum gap size, distance between seed and extension, and allowed curvature were set by the user. Motors were identified using a difference of Gaussian filtered images and a user-set threshold with user defined minimum distance to MT seed or extension. A MT binding event at the plus or minus end was considered when the motor binding event was within three pixels or 0.3 μm from the MT end. A user driven interface allowed editing of all segmented data points overlain onto original images for verification. Results from each image were output to Excel files where they were compiled and then imported into Prism 9 for graphing and statistical analysis. In addition, output figures were generated for each image that displayed the MTs from each MT class (seed only, seed with two extensions (2-sided), seed with one extension (1-sided), and extension only) with the corresponding positioned motor binding events as identified and as linearized representations of polymer. A second MATLAB GUI was developed to compile the data from each experimental condition from independent experiments into single output Excel files and linear representations of seed, extension, and motor position. Data presented is from three independent experiments.

The percentage of MTs bound with GFP-Kif18B motor ± a four-fold excess of importins or EB1 was determined by dividing the number of 2-sided MTs that had at least one motor binding event by the total number of 2-sided MTs in that condition for each experiment, averaged, and graphed as the mean ± SEM. An unpaired Student’s *t*-test was used to determine significant differences from GFP-Kif18B alone. The percentage of MTs with motor binding events at the distal plus or minus ends of the 2-sided MTs in each condition was determined by dividing the number of events at either end by the number of bound MTs, averaged, and graphed as the mean ± SEM. The criteria for determining whether a minus end with a motor binding event might be a short plus end, was considered when the plus-end MT length was shorter than 90% of control GFP minus-end MT length or 2.7 um. A multiple unpaired *t*-test was used to determine significance compared to GFP-Kif18B alone. The number of motor binding events per MT was plotted for each condition with the median ± interquartile range (IQR). The distribution of binding events was lognormal based on the Normality and Lognormality Test in GraphPad Prism 9. ANOVAs using the Kruskal-Wallis test with Dunn’s multiple comparisons were performed for significance from WT GFP-Kif18B alone. MT lengths of the 2-sided MTs, the 1-sided MTs, the seeds, and the plus- and minus-end extensions had non-normal distributions as indicated by the Normality and Lognormality Tests in Prism. To best visualize these distributions, the lengths were graphed on a log scale with the median ± IQR. Significant differences in their median lengths were determined using the Kruskal-Wallis test with Dunn’s multiple comparisons test. Significance was considered when p < 0.05 for statistical tests.

### Single Motor Motility Experiments

MTs were polymerized by incubating 5 µM tubulin (4% HyLite 647 tubulin (Cytoskeleton #TL670M), ∼1% biotin tubulin (Cytoskeleton #T333P), 95% unlabeled cycled bovine tubulin) in BD with 0.5 mM GMPCPP for 30 min, sedimented, and the MT pellet resuspended in BD. MT concentration was determined using an extinction coefficient of 115,000 M^-1^ cm^-1^ for tubulin dimer. Coverslips (24×30 mm and 22×22 mm #1.5) were washed and treated with VECTABOND® Reagent (Vector Laboratories, Newark, CA) and then coated with 25% polyethylene glycol (PEG), 0.7% biotin-polyethylene glycol, rinsed, dried with nitrogen, and stored under vacuum as previously described (Ems-McClung *et al*., 2013). Flow chambers were constructed fresh for each experiment from a sandwich of a 24×30 mm and a 22×22 mm PEG coated coverslip with double-stick tape to create three equally spaced chambers parallel to the long edge of the coverslip affixed to a glass slide with double-stick tape. Each chamber held ∼10 uL solution. Chambers were treated with 5% Pluronic F-127 in BD for 3 min, rinsed with Block (BD, 50 mM KCl, 2 mM MgATP, 1 mg/ml casein), and treated with 0.05 mg/ml Neutravidin in Block for 3 min. Chambers were then rinsed with Block, and 0.15 µM GMPCPP-stabilized HyLite 647/biotin-labeled MTs diluted in Block were introduced and incubated for 3 min. All incubations were done with the slide upside down. The chamber with immobilized MTs was rinsed with Block + OSM (0.32 mg/ml glucose oxidase, 55 µg/ml catalase, 5 mM DTT, 25 mM glucose, 50 mM KCl, BD), and then 0.05 nM GFP-Kif18B ± 0.2 nM importin α-His_6_/His_6_-S-importin β or His_6_-importin α-mCherry/His_6_-S-importin β in Block + OSM was introduced and imaged immediately using total internal reflection fluorescence (TIRF) microscopy on a Nikon Eclipse Ti2 microscope equipped with a Nikon 100× Apo TIRF NA 1.49 oil objective, Coherent OBIS lasers (488, 561, 637 nm), and an Andor EMCCD iXon3 camera controlled by Micro-Manager 2.0.

For imaging, single plane images of MTs were taken at the beginning and end of each movie using the 637 nm laser at 1 mW power for 100 ms. GFP-Kif18B motility was imaged with the 488 nm laser at 1 mW power with 50 ms exposures for 1200 frames at 11.6 frames per second (fps). GFP-Kif18B and importin α-mCherry were imaged sequentially with GFP-Kif18B being imaged at 1 mW power with 50 ms exposures and importin α-mCherry being imaged with 5 mW power and 50 ms exposures for 300 frames at 0.76 fps. Data was collected from two or three fields of view from each imaging condition (11.6 fps or 0.76 fps) for each protein condition (GFP-Kif18B or GFP-Kif18B + importin α/β) per experiment. Four independent experiments were analyzed.

Color combined images of the Cy5 MT channel and the maximum projected GFP or mCherry channel were generated in FIJI (NIH) and MTs identified with binding events based on colocalization of the MT and GFP or mCherry signals. GFP-Kif18B or importin α-mCherry movies were assembled with the MT image as the first image followed by the GFP or mCherry image series. Ten to 15 kymographs were generated per movie by drawing a line across the MT and extending slightly past the ends that were identified in the color combined images using the Multi Kymograph macro in FIJI. Each kymograph was rotated 90° to the left, the length of the MT measured in the first frame, and the binding events measured using the line tool. Events were scored if they persisted for at least three timeframes and were scored as either being diffusive or moving. A diffusive event was scored if its displacement did not change or changed less than three pixels in a discontinuous manner. A moving or directed motility event was scored if its displacement was in only one direction and persisted for three or more frames. Kymograph results were exported to Excel. The mean percentages of diffusive and moving events were determined for each condition per experiment in Excel and graphed as the mean ± SEM in Prism 9. The time an event persisted was determined by dividing the number of timeframes by the frame rate (11.6 fps). The MT length and the run length of moving events was determined by multiplying the height times 0.160 µm/pixel. Velocity was calculated from the run length divided by the time, and the on-rate was calculated by dividing the number of landing events starting and stopping within the timeframe of the movie by the length of the MT in microns and the length of the movie in seconds. An event was scored at the plus end if it was within 3 pixels of the plus end of the measured length of the MT. An event was categorized as a lattice event if it was further than 3 pixels from either MT end. The polarity of the MT was determined based on the direction of moving GFP-Kif18B molecules.

For data analysis and graphical representation, on-rates, velocities, run lengths, and dwell times were assembled in Prism 9. The distribution of the on-rates (*k*_on_) and velocities were lognormal based on normality tests; thus, Mann-Whitney U tests were performed. The individual values are plotted on a log_10_ scale to best present the data with, and the geometric means and with the geometric SD factor indicated. The average run length of moving GFP-Kif18B molecules was plotted as a histogram and fit to a single exponential decay model, and the mean run length, tau (τ), was determined from the inverse of the rate constant. Significant differences between the run lengths of GFP-Kif18B ± importins were determined by an extra sum-of-squares F test of the rate constant. The mean run length with 95% confidence intervals (CI) is reported. The GFP-Kif18B dwell times of diffusive events on the lattice or plus ends without or with importin α/β were plotted as a histogram and fit to a single exponential decay model to determine the observed off-rate, *k*_Obs_. The corrected mean dwell time, τ, was determined from the inverse of the corrected off-rate, *k*_off_. The observed off-rates were corrected for photobleaching by subtracting the photobleaching rate constant, *k*_B_, to determine *k*_off_. The photobleaching rate constant was determined from two-step photobleaching events that occurred on the MTs as well as in the background as described for the lattice and plus end dwell times. The rate constant for photobleaching equaled 0.0517 s^-1^. Significant differences between the dwell times of GFP-Kif18B ± importins were determined by an extra sum-of-squares F test of the off-rate. The corrected mean dwell times with 95% confidence intervals (CI) are reported and graphed as a histogram with the exponential fit. Significance was considered when p<0.05 such that *, p<0.05; **, p<0.01; ***, p<0.001; ****, p<0.0001.

## Acknowledgements

This work was supported by NIH R35GM122482 to CEW, NSF MCB2128166 to CEW, and NSF MCB1615907 to SLS. Support for the OMX 3D-SIM microscope was provided by NIH S10 OD024988-01. We thank Anjaly Prasannajith for help in making the Kif18B lentiviral constructs, and Puck Ohi and Jordon Winn for comments on the manuscript. We thank Yan Yu for allowing us to use her TIRF imaging system and to Seonik Lee for assistance in using the TIRF system. We are indebted to Jim Powers and Andras Kun in the IU-Light Microscopy Imaging Center for their assistance with the OMX imaging and Amber English from IMARIS for assistance in the reconstructions and data analysis. The IUB-LMIC is supported in part by the Office of the Vice Provost for Research.

## Supplemental Material

### Supplemental Figures

**Figure S1.**
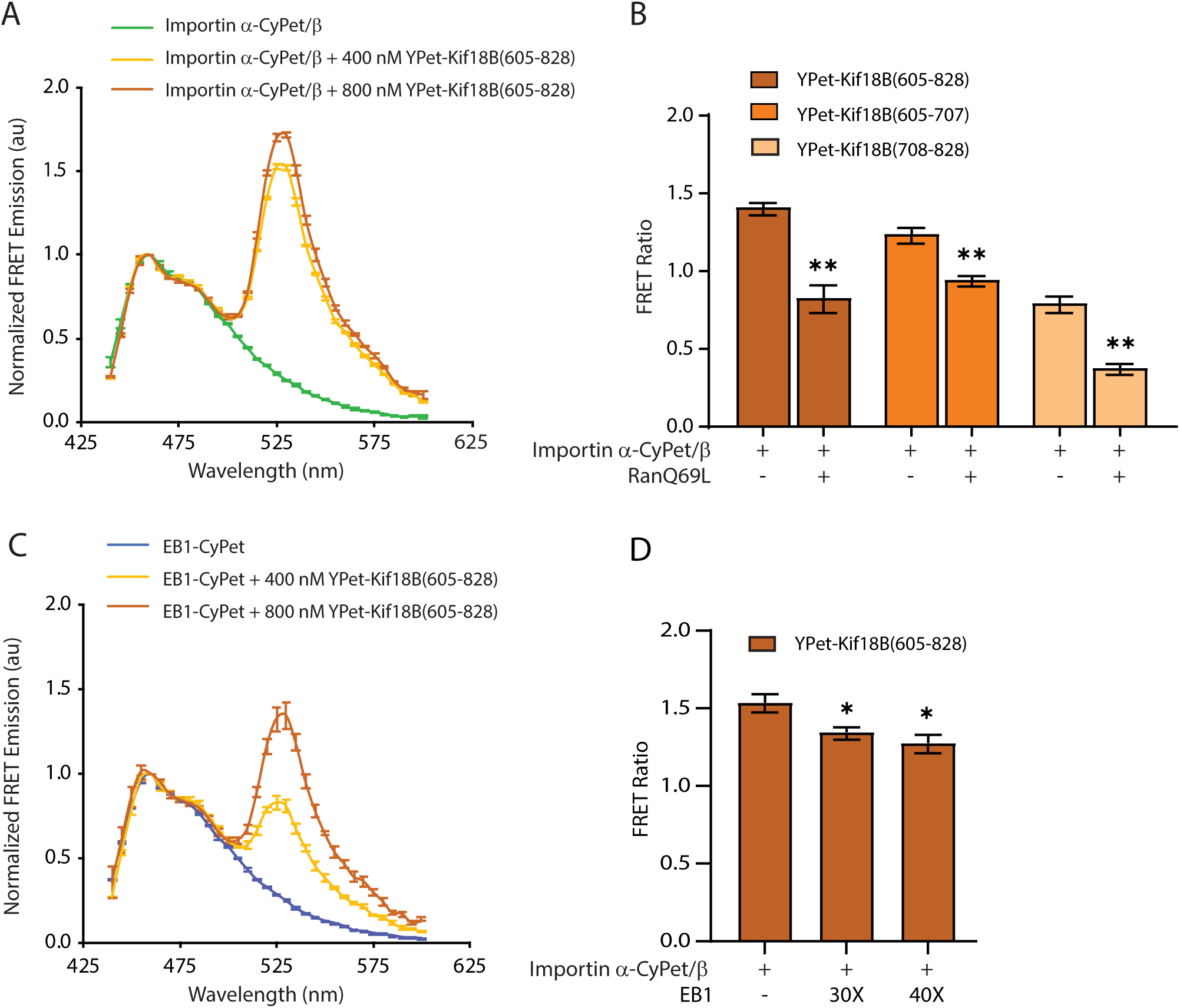
Kif18B undergoes intermolecular FRET with importin a/β or EB1 in a dose dependent manner. Reactions containing 100 nM importin α-CyPet and 400 nM importin β (A) were incubated with 400 or 800 nM YPet-Kif18B(605-828), excited at 405 nm, and their spectral emissions recorded from 440 to 600 nm in 5 nm steps. (B) Reactions containing 100 nM importin α-CyPet and 400 nM importin β with 400 nM YPet-Kif18B tail proteins were measured for FRET as in (A) and then incubated with 10 μM RanQ69L for 10 min and their FRET re-measured from three independent experiments. The mean FRET ratio ± SEM before and after RanQ69L addition are plotted. (C) 100 nM of EB1-CyPet (B) was incubated with 400 or 800 nM of YPet-Kif18B(605-828) and the FRET measured as in (A). (A and C) The average normalized spectral emissions from three independent experiments are graphed as the mean ± SEM. (D) Reactions containing 100 nM importin α-CyPet and 400 nM importin β with 600 nM YPet-Kif18B(605-828) in the absence or presence of 3 µM (30X) or 4 µM 6His-EB1 (40X) were measured for FRET in three independent experiments as in (A). The mean FRET ratio ± SEM is plotted. *, p<0.05; **, p<0.01.

**Figure S2.**
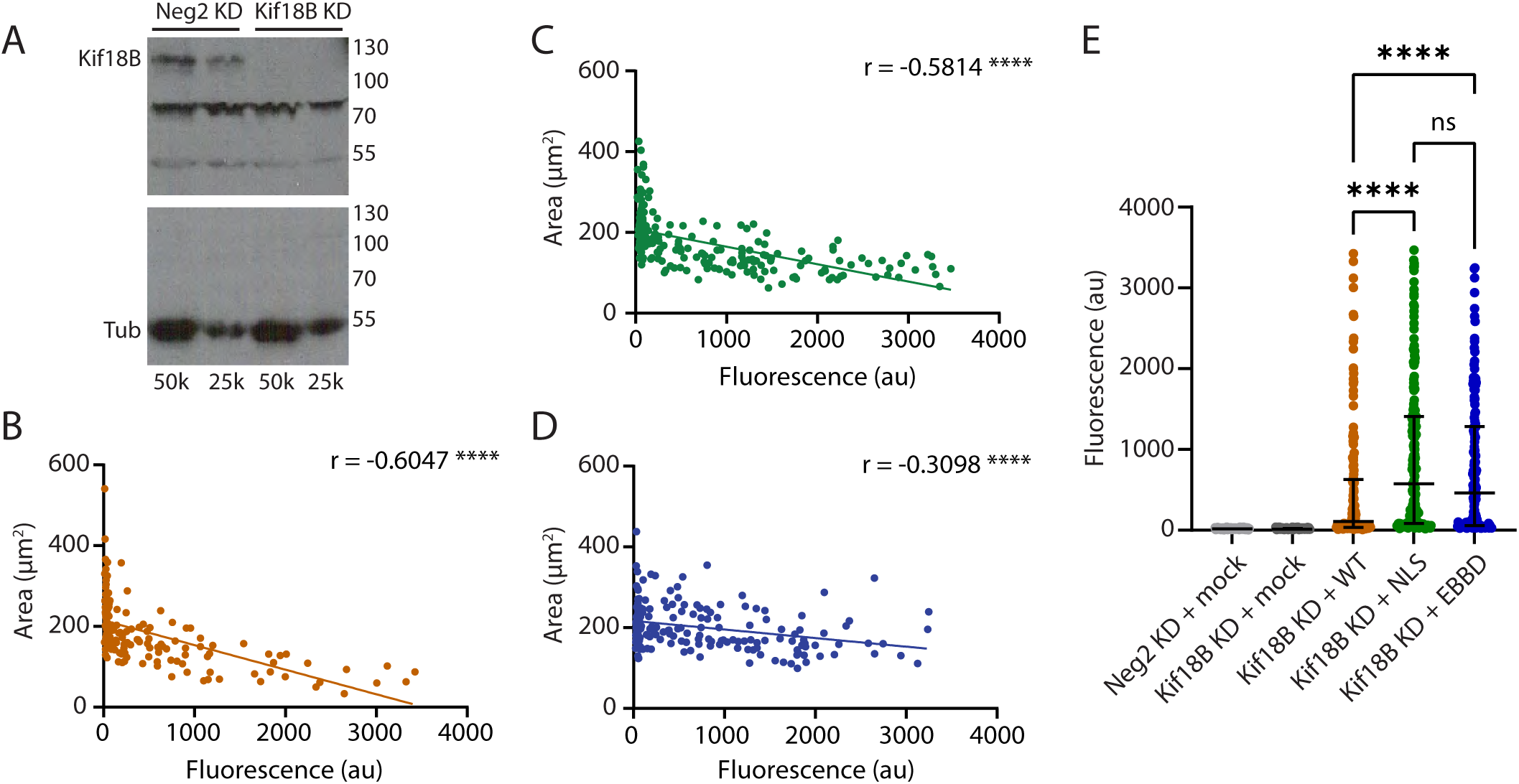
Analysis of GFP expression levels versus aster area. (A) Western blots of control Neg2 knockdown cells (Neg2 KD) or Kif18B knockdown cells (Kif18B KD) probed with an anti-Kif18B antibody (top) or α-tubulin antibody as a loading control (bottom). The number of cells loaded in each lane is indicated at the bottom of the Western. (B-D) Correlation between the aster area vs. GFP fluorescence intensity for all cells measured in Figure 4 expressing WT GFP-Kif18B (B), GFP-Kif18B NLS (C) or GFP-Kif18B EBBD (D). (E) Dot plots showing the individual expression level of GFP in all cells measured for analysis of aster area. This data is before binning for equivalent expression levels between constructs as shown in Figure 4B. ns, not significant; ****, p<0.0001.

**Figure S3.**
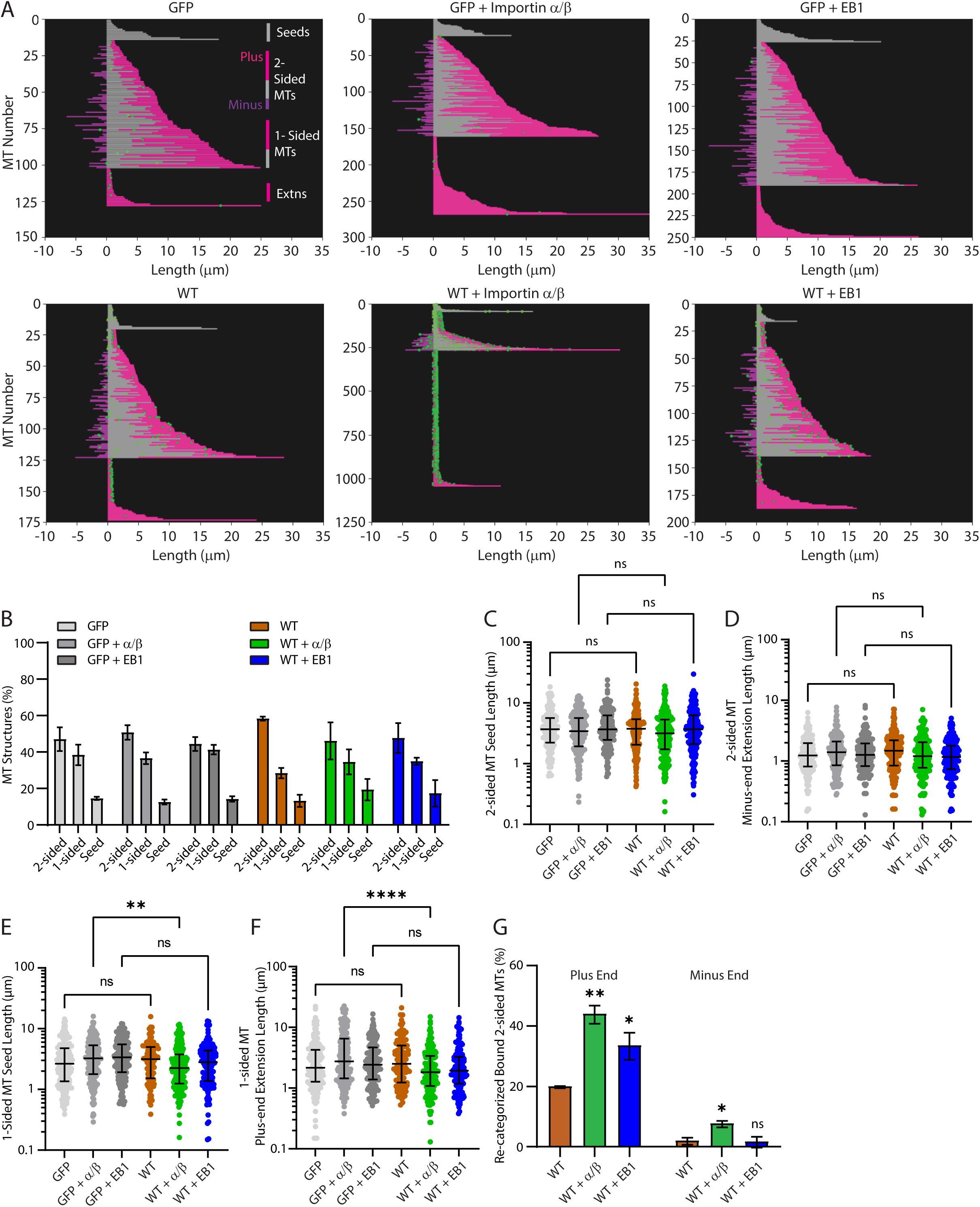
EB1 has increased distal MT plus-end binding after re-categorization of MT lengths. (A) Complete data sets for each condition of a representative experiment analyzed by the custom-built MATLAB algorithm for GFP-Kif18B MT binding and length determination represented in Figure 7 where seeds are gray, plus-end extensions are magenta, and minus-end extensions are purple. (B) The percentage of MT structures that were 2-sided, 1-sided, or seed only plotted as mean ± SEM. (C) The lengths of the 2-sided MT seeds (C), 2-sided minus-end extensions (D), 1-sided MT seeds (E) or 1-sided plus-end MT extensions (F) were plotted for each MT on a log scale to better represent the lognormal distribution with the median ± interquartile range. A Kruskal-Wallis ANOVA with Dunn’s multiple comparisons test was performed to compare the medians. (G) MTs with minus-end MT binding events were re-categorized as plus-end MT binding events if the longer extension was within 90% of the length of control GFP minus-end MT lengths or 2.7 µm. The resulting adjusted percentage of plus-end and minus-end MT binding is plotted from the three independent experiments as the mean ± SEM. A multiple *t*-test was performed to compare the means. ns, not significant; *, p<0.05; **, p<0.001; ****, p<0.0001.

**Figure S4.**
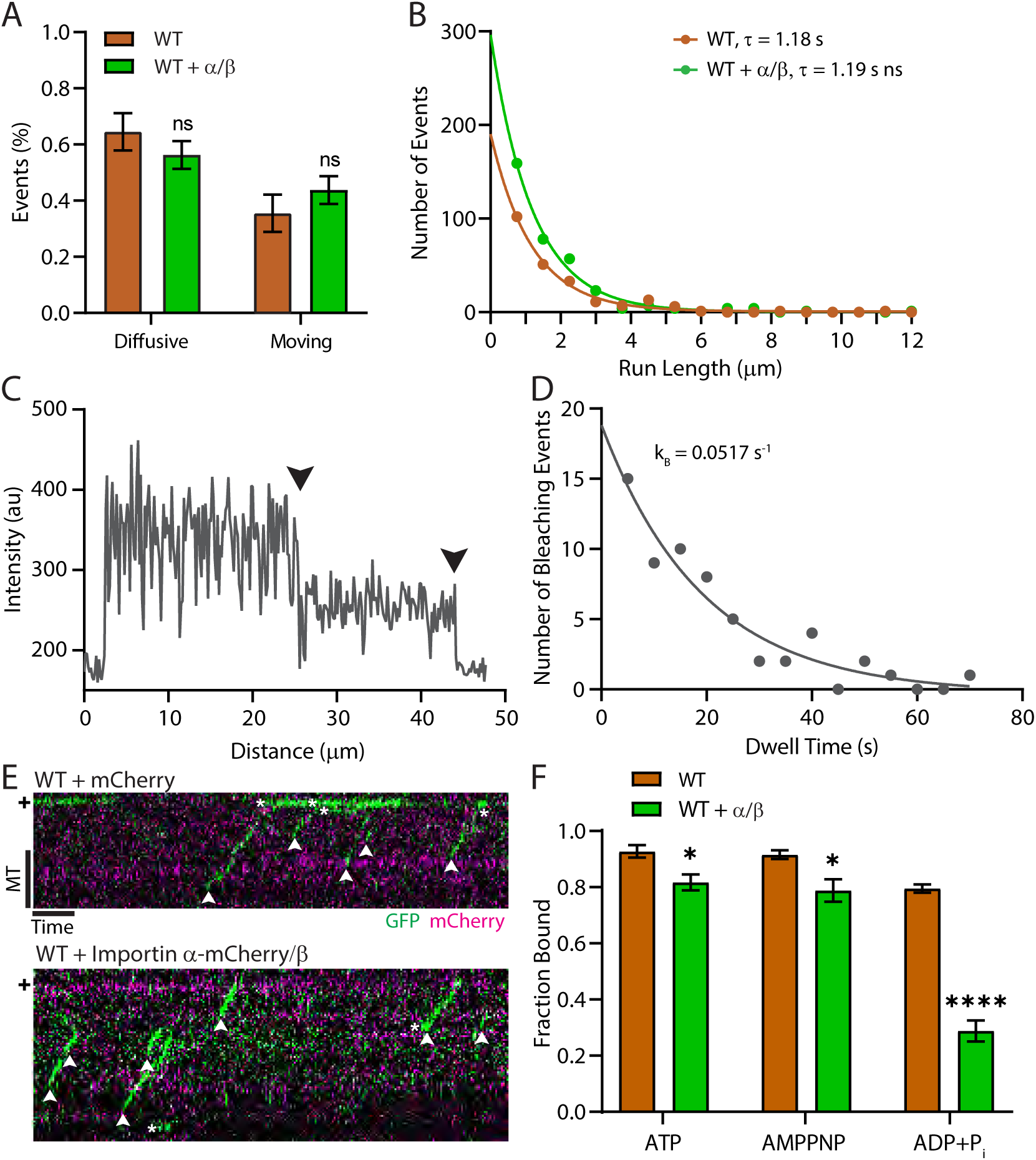
**I**mportin α/β do not affect the diffusive or directed motility of Kif18B. (A) The fraction of diffusive and directed moving events were calculated from four independent experiments and graphed as the mean ± SEM. An unpaired Student’s *t*-test was performed to compare the means. (B) The run lengths of moving GFP-Kif18B ± **i**mportin α/β are graphed as a histogram and fit to a single exponential decay model to determine the mean run length, τ. The mean run lengths were compared using an extra sum-of-squares F test on the rate constant from the single exponential decay curve and are indicated. (C and D) The photobleaching correction factor, *k*_B_, for the off-rates of GFP-Kif18B were determined from two-step bleaching events (C) on MTs and events non-specifically bound to the coverslip as described in Figure 8D for lattice dwell times (D). (E) Representative kymographs of sequential imaging of GFP-Kif18B + mCherry (top) or GFP-Kif18B + importin α/β (bottom) depicted as described in Figure 8A. MT scale bar, 5 µm and time scale bar, 30 s. (F) Fractional MT binding of 200 nM GFP-Kif18B ± 800 nM importin α/β in 2 mM MgATP, 5 mM MgAMPPNP, 2 mM MgADP + 20 mM P_i_ with 1 µM doubly stabilized MTs are graphed as the mean ± SEM from at least three independent experiments. An unpaired Student’s *t*-test was performed to compare the means. ns, not significant; *, p<0.05; ****, p<0.0001.

**Video 1.** Importin α/β increase the off-rate and lattice dwell time of Kif18B. Movies of single molecules of GFP-Kif18B (WT, green) and GFP-Kif18B + importins (WT + Importin α/β, green) showing diffusing (*) or moving (^) molecules along a MT (magenta). Scale bar, 2 µm.

**Video 2.** Importin α-mCherry/importin β do not associate with GFP-Kif18B during diffusive or moving events. Movies of sequential imaging of single molecules of GFP-Kif18B (green) + mCherry (magenta) or GFP-Kif18B (green) + importin α-mCherry/importin β (magenta) diffusing (*) or moving along a MT (^, gray). Scale bar, 2 µm.

### Supplemental Tables

**Table S1.**
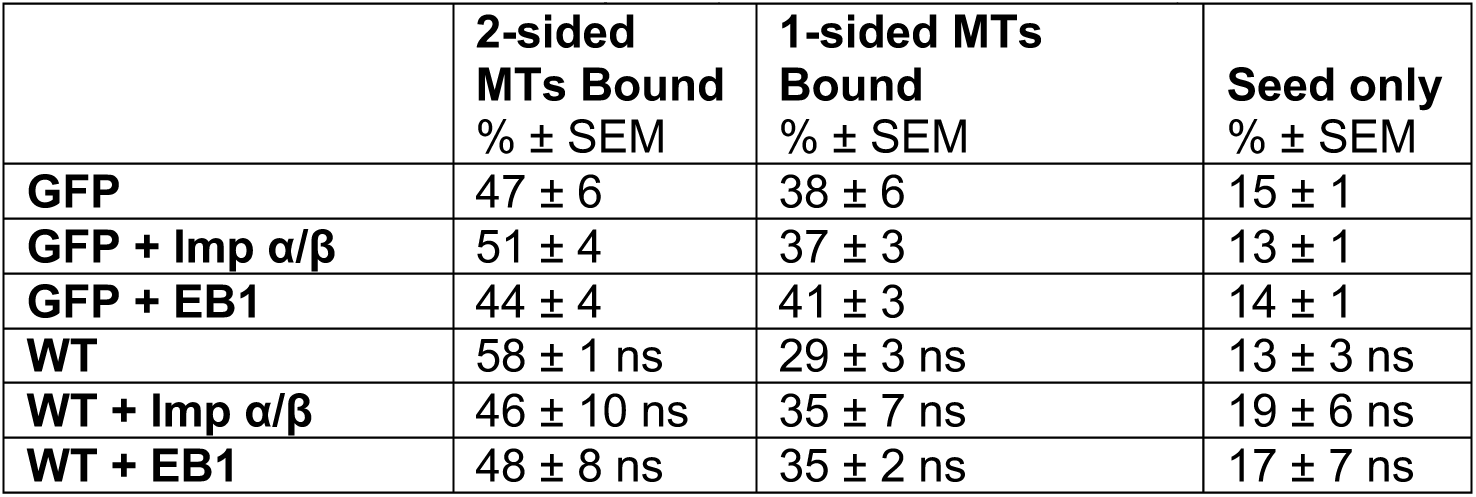
Percent MT Binding in Dynamic MT Seed Assays The mean percentage of 2-sided MTs, 1-sided MTs, and Seed only MT structures from three independent experiments are reported with the standard error of the mean (SEM) and graphed in Figure S3B.

| | 2-sided MTs Bound<br>% ± SEM | 1-sided MTs Bound<br>% ± SEM | Seed only<br>% ± SEM |
| --- | --- | --- | --- |
| **GFP** | 47 ± 6 | 38 ± 6 | 15 ± 1 |
| **GFP + Imp $\alpha/\beta$** | 51 ± 4 | 37 ± 3 | 13 ± 1 |
| **GFP + EB1** | 44 ± 4 | 41 ± 3 | 14 ± 1 |
| **WT** | 58 ± 1 ns | 29 ± 3 ns | 13 ± 3 ns |
| **WT + Imp $\alpha/\beta$** | 46 ± 10 ns | 35 ± 7 ns | 19 ± 6 ns |
| **WT + EB1** | 48 ± 8 ns | 35 ± 2 ns | 17 ± 7 ns |

**Table S2.** Location of MT Binding Events in Dynamic MT Seed Assays The mean percentages of 2-sided or 1-sided MTs bound, plus-end bound, or minus-end bound with GFP-Kif18B ± importins or EB1 are reported with the SEM and graphed in Figure 7, B and C and E and F. The median number of binding events per 2-sided (n=95-174) or 1-sided (n=35-116) MTs relative to WT GFP-Kif18B and interquartile range (IQR) are reported and graphed in Figure 7D (2-sided) or Figure 7G (1-sided) from three independent experiments. ns, not significant; *, p<0.05; **, p<0.01; ***, p<0.001.

| | MTs Bound<br>% ± SEM | Plus-End Bound<br>% ± SEM | Minus-End Bound<br>% ± SEM | Number of Binding Events per MT<br>Median (IQR) | Number |
| --- | --- | --- | --- | --- | --- |
| 2-sided MTs |  |  |  |  |  |
| **WT** | 46 ± 6 | 17 ± 2 | 5 ± 1 | 1 (1 to 2) | 95 |
| **WT + Imp $\alpha/\beta$** | 73 ± 7 * | 33 ± 2 ** | 19 ± 5 * | 2 (1 to 3) *** | 98 |
| **WT + EB1** | 50 ± 6 ns | 23 ± 3 ns | 13 ± 2 * | 1 (1 to 2) ns | 174 |
| 1-sided MTs |  |  |  |  |  |
| **WT** | 36 ± 5 | 6 ± 5 | 3 ± 0.4 | 1 (1 to 2) | 35 |
| **WT + Imp $\alpha/\beta$** | 60 ± 3 * | 20 ± 1 * | 8 ± 2 * | 2 (1 to 2) * | 47 |
| **WT + EB1** | 34 ± 8 ns | 7 ± 2 ns | 5 ± 4 ns | 1 (1 to 2) ns | 116 |

**Table S3.** Lengths of MTs in Dynamic MT Seed Assays The MT lengths, plus-end extension lengths, seed lengths, and minus-end extension lengths of 2-sided or 1-sided MTs with WT GFP-Kif18B in the absence or presence of importin α/β or EB1 are reported as the median and interquartile range (IQR) and graphed in Figures, 7H-I and S3C-D (2-sided) or Figures, 7J and S3E-F (1-sided) from three independent experiments. ns, not significant; *, p<0.05; **, p<0.01; ****, p<0.0001.

| 2-sided MTs | MT Length<br>μm (IQR) | Plus-End<br>Length<br>μm (IQR) | Seed Length<br>μm (IQR) | Minus-End<br>Length<br>μm (IQR) | Number |
| --- | --- | --- | --- | --- | --- |
| **GFP** | 9.5 (7.2 to 13.9) | 3.6 (2.5 to 6.6) | 3.7 (2.2 to 5.7) | 1.2 (0.8 to 2.0) | 175 |
| **GFP + Imp $\alpha/\beta$** | 10.6 (7.6 to 14.9) | 4.6 (2.7 to 7.8) | 3.4 (1.9 to 5.6) | 1.4 (0.8 to 2.1) | 235 |
| **GFP + EB1** | 10.2(7.9 to 14.1) | 4.2 (2.2 to 6.6) | 3.7 (2.5 to 6.2) | 1.3 (0.8 to 1.9) | 192 |
| **WT** | 9.8 (7.2 to 13.8)<br>ns | 3.7 (2.5 to 6.5)<br>ns | 3.7 (2.1 to 5.4)<br>ns | 1.5 (0.8 to 2.2)<br>ns | 208 |
| **WT + Imp $\alpha/\beta$** | 8.6 (5.9 to 11.8)<br>**** | 3.0 (2.0 to 5.3)<br>**** | 3.2 (1.7 to 5.3)<br>ns | 1.2 (0.8 to 2.0)<br>ns | 248 |
| **WT + EB1** | 9.5 (6.8 to 12.8)<br>ns | 3.1 (2.0 to 5.0)<br>* | 3.7 (2.1 to 6.2)<br>ns | 1.2 (0.7 to 1.8)<br>ns | 197 |
| 1-sided MTs |  |  |  |  |  |
| **GFP** | 5.8 (3.6 to 8.4) | 2.2 (1.3 to 4.2) | 2.6 (1.4 to 4.8) | NA | 165 |
| **GFP + Imp $\alpha/\beta$** | 7.7 (4.7 to 10.7) | 2.7 (1.4 to 6.5) | 3.2 (1.8 to 5.3) | NA | 175 |
| **GFP + EB1** | 6.7 (4.1 to 10.0) | 2.4 (1.4 to 4.7) | 3.4 (1.9 to 5.5) | NA | 182 |
| **WT** | 6.4 (4.0 to 9.9)<br>ns | 2.5 (1.2 to 5.0)<br>ns | 3.1 (1.5 to 5.0)<br>ns | NA | 102 |
| **WT + Imp $\alpha/\beta$** | 4.6 (2.9 to 7.1)<br>**** | 1.8 (1.1 to 3.4)<br>**** | 2.2 (1.2 to 3.8)<br>** | NA | 198 |
| **WT + EB1** | 5.6 (3.3 to 7.2)<br>* | 1.9 (1.2 to 3.3)<br>ns | 2.8 (1.4 to 4.4)<br>ns | NA | 149 |

**Table S4.** Motility Events in Single Motor Assays The mean percentage of diffusive or directed moving events in reactions with WT GFP-Kif18B ± importin α/β are reported with the SEM from four independent experiments and graphed in Figure S4A. The on-rates (kon) and velocities of WT GFP-Kif18B ± importin α/β are reported as the geometric mean with the geometric SD factor and graphed in Figure 8, B and C. The mean run length, τ, of GFP-Kif18B ± importin α/β was determined from the inverse of the rate constant and is reported with 95% confidence intervals (CI) and graphed as a histogram with the exponential fit in Figure S4B. Data is from four independent experiments. ns, not significant; *, p<0.05.

| | Diffusive<br>% ± SEM | Moving<br>% ± SEM | On-rate, $k_{on}$<br>μm·s<sup>-1</sup> (Geo<br>SD) | Number | Velocity<br>μm s<sup>-1</sup> (Geo<br>SD) | Run Length, $\tau$<br>μm (95% CI) | Number |
| --- | --- | --- | --- | --- | --- | --- | --- |
| **WT** | 64 ± 7 | 36 ± 7 | 0.26 (2.95) | 139 | 0.26 (1.76) | 1.18<br>(1.01 to 1.37) | 229 |
| **WT + Imp $\alpha/\beta$** | 56 ± 5 | 44 ± 5 | 0.35 (2.92)<br>* | 133 | 0.25 (1.74)<br>ns | 1.19<br>(1.04 to 1.36)<br>ns | 346 |

**Table S5.** On and Off Rates in Single Motor Assays The mean dwell time, τ, of diffusive events on the lattice or plus ends of GFP-Kif18B ± importin α/β were determined from the inverse of the off-rate, *k*_off_, that was corrected for photobleaching. The corrected mean dwell times with 95% confidence intervals (CI) are reported and graphed as a histogram with the exponential fit in Figure 8, D and E from four independent experiments. ns, not significant; ***, p<0.001.

|  | **Lattice Off-rate, $k_{off}$**<br>s<sup>-1</sup> (95% CI) | **Lattice Dwell Time, $\tau$**<br>s (95% CI) | **Number** | **Plus-End Off-rate, $k_{off}$**<br>s<sup>-1</sup> (95% CI) | **Plus-End Dwell Time, $\tau$**<br>s (95% CI) | **Number** |
| --- | --- | --- | --- | --- | --- | --- |
| **WT** | 0.56<br>(0.50 to 0.63) | 1.77<br>(1.58 to 2.00) | 278 | 0.18<br>(0.13 to 0.25) | 5.56<br>(4.04 to 7.78) | 47 |
| **WT + Imp $\alpha/\beta$** | 0.43<br>(0.40 to 0.47)<br>*** | 2.30<br>(2.14 to 2.47) | 336 | 0.19<br>(0.11 to 0.29)<br>ns | 5.39<br>(3.45 to 8.71) | 70 |

**Table S6.** Recategorization of MT End Definition in Dynamic MT Seed Assays The mean percentage of re-categorized 2-sided MTs, plus-end extensions, and minus-end extensions bound with GFP-Kif18B ± importins or EB1 are reported with the SEM and graphed in Figure S3G from three independent experiments. ns, not significant; *, p<0.05; **, p<0.01.

| **Re-categorized 2-sided MTs** | **Plus-End Bound**<br>% ± SEM | **Minus-End Bound**<br>% ± SEM |
| --- | --- | --- |
| **WT** | 20 ± 0.3 | 2 ± 1 |
| **WT + Imp $\alpha/\beta$** | 44 ± 3 ** | 8 ± 1 * |
| **WT + EB1** | 34 ± 5 * | 2 ± 2 ns |

**Table S7.**
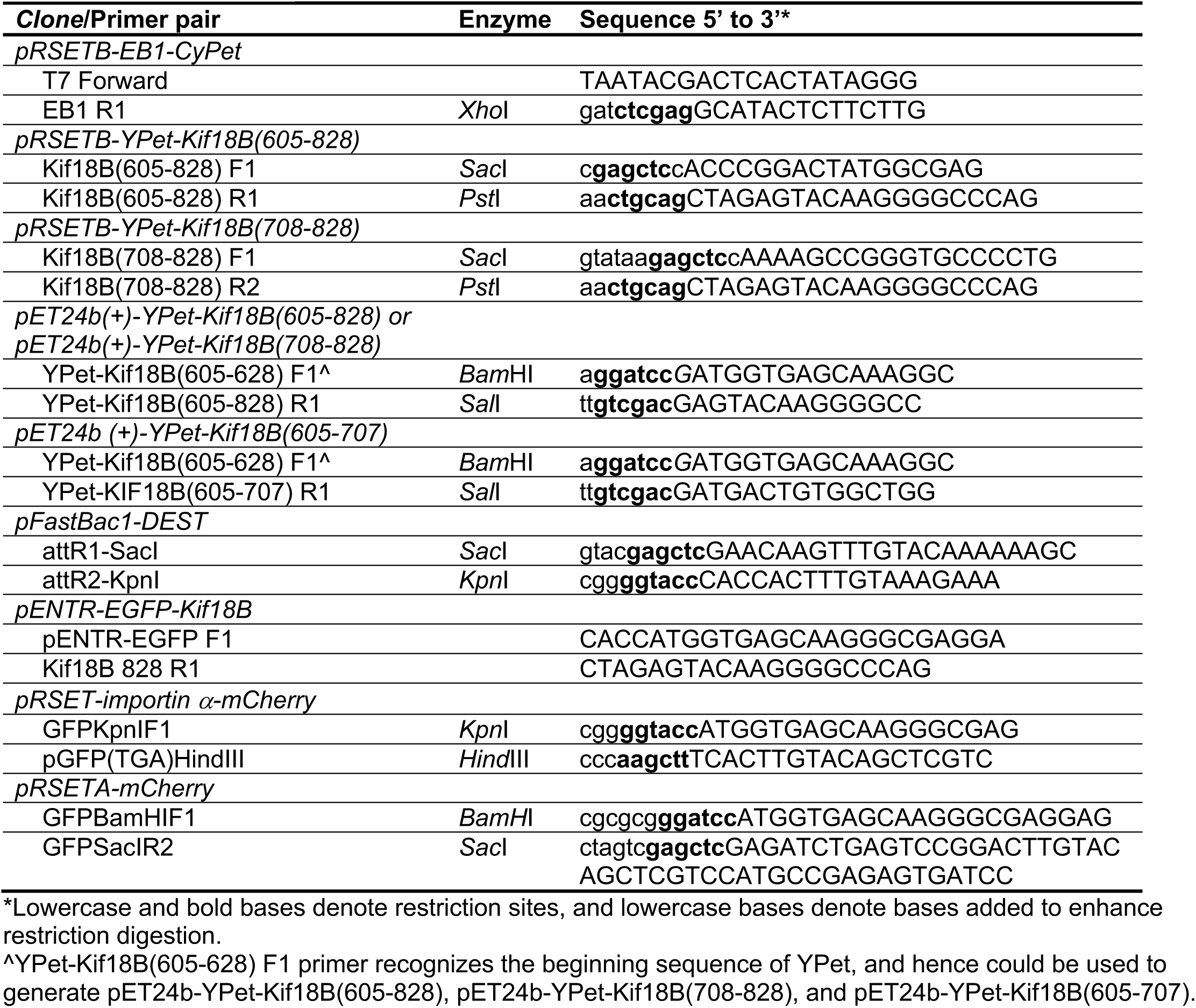
Cloning primers

**Table S8.**
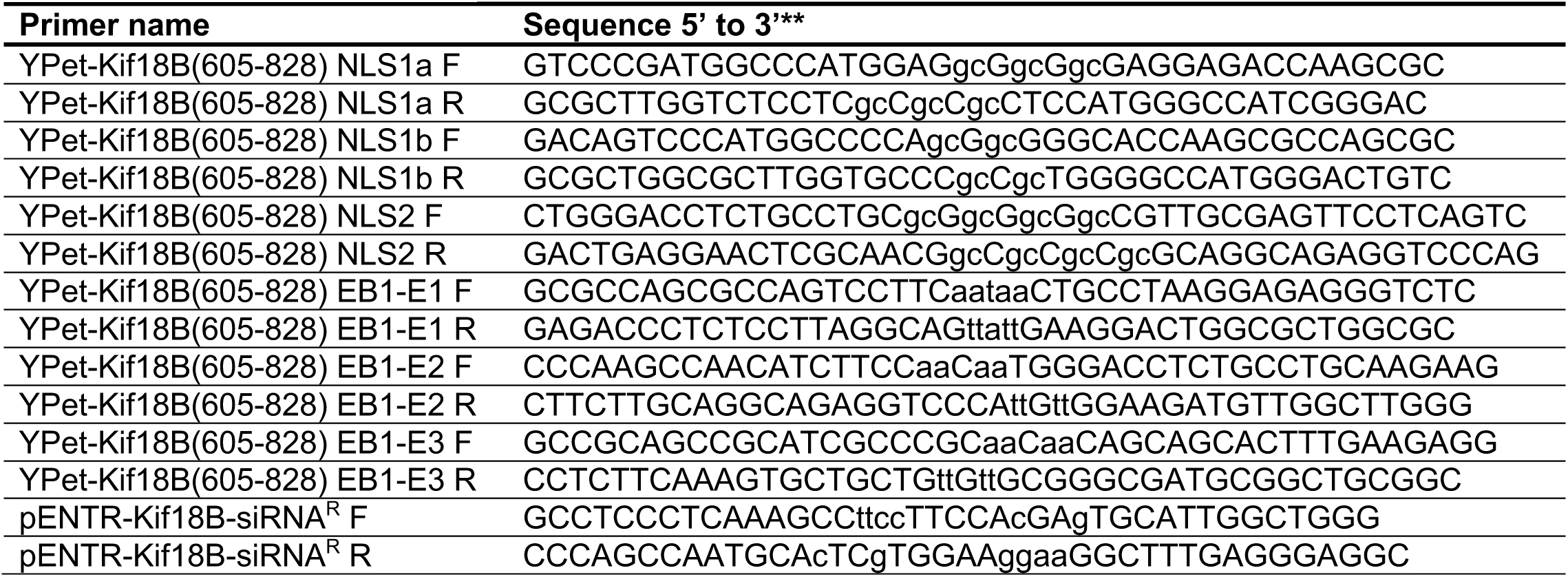
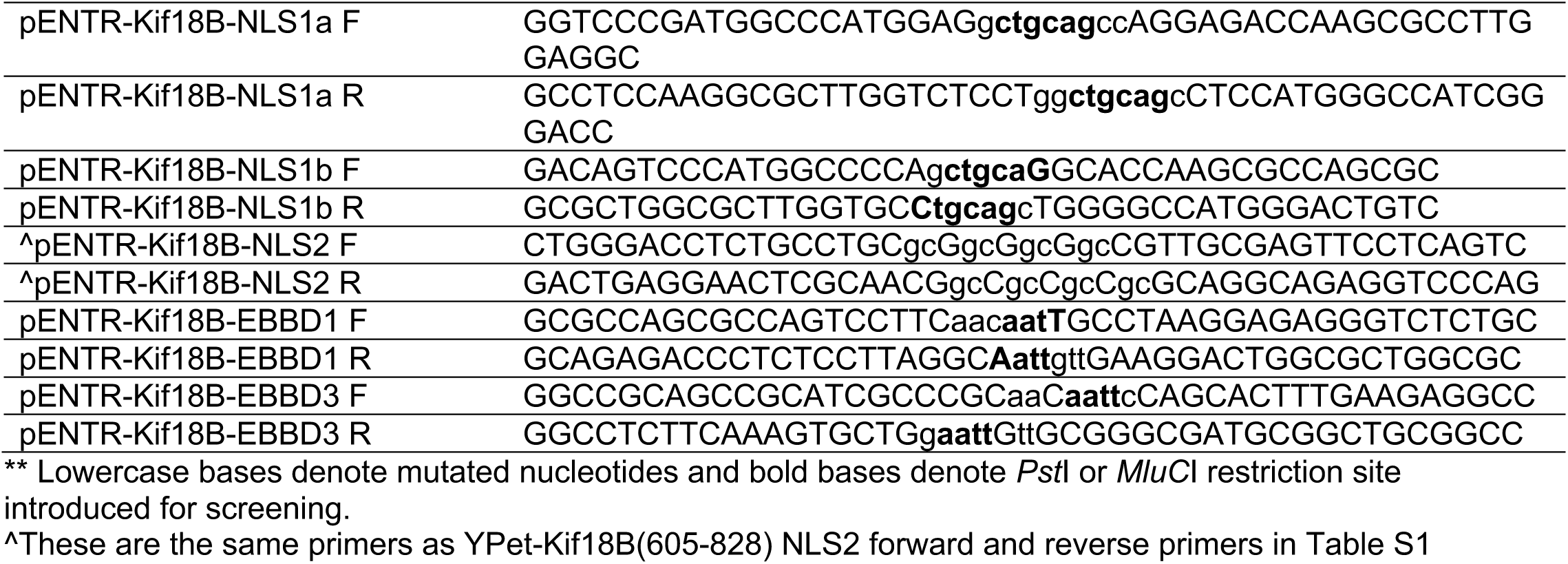
Mutagenesis primers

## Notes

### Competing Interest Statement

The authors have declared no competing interest.

### Summary of Updates

The revision adds new data looking at the mechanism by which Kif18B enhances MT destabilization activity as well as provides additional supporting data.

